# Reconstructing essential active zone functions within a synapse

**DOI:** 10.1101/2021.11.21.469466

**Authors:** Chao Tan, Shan Shan H. Wang, Giovanni de Nola, Pascal S. Kaeser

## Abstract

Active zones are molecular machines that control neurotransmitter release through synaptic vesicle docking and priming, and through coupling of these vesicles to Ca^2+^ entry. The complexity of active zone machinery has made it challenging to determine which mechanisms drive these roles in release. Here, we induce RIM+ELKS knockout to eliminate active zone scaffolding networks, and then reconstruct each active zone function. Re-expression of RIM1-Zn fingers positioned Munc13 on undocked vesicles and rendered them release-competent. Reconstitution of release-triggering required docking of these vesicles to Ca^2+^ channels. Fusing RIM1-Zn to Ca_V_β4-subunits sufficed to restore docking, priming and release-triggering without reinstating active zone scaffolds. Hence, exocytotic activities of the 80 kDa Ca_V_β4-Zn fusion protein bypassed the need for megadalton-sized secretory machines. These data define key mechanisms of active zone function, establish that fusion competence and docking are mechanistically separable, and reveal that active zone scaffolding networks are not required for release.

## Introduction

Essential insight into the functioning of synaptic exocytotic machinery has come from rebuilding the fusion process in vitro. Reconstitution assays have revealed that the minimal machinery for Ca^2+^-triggered exocytosis consists of SNARE proteins, Munc18, Munc13 and synaptotagmin (Hu et al., 2003; Ma et al., 2013; Tucker et al., 2004). However, fusion speed in these assays is orders of magnitude slower than at synapses, and spatial precision of exocytosis with its exact targeting towards receptors on target cells cannot be studied in these in vitro systems. These functions are carried out by the active zone, a molecular machine that is attached to the presynaptic plasma membrane and is composed of many megadalton-sized protein assemblies (Emperador-Melero and Kaeser, 2020; Südhof, 2012).

Central functions of the active zone are the generation of releasable vesicles and the positioning of these vesicles close the Ca^2+^ channels for rapid fusion-triggering (Augustin et al., 1999; Biederer et al., 2017; Deng et al., 2011; Imig et al., 2014; Kaeser et al., 2011; Liu et al., 2011). Active zones are composed of families of scaffolding proteins including RIM, ELKS, Munc13, RIM-BP, Bassoon/Piccolo and Liprin-α (Südhof, 2012). Each of these proteins is encoded by multiple genes and the individual proteins are large, ranging from 125 to 420 kDa, forming complex protein networks. Mechanisms for their assembly are not well understood, but recent studies suggest the involvement of liquid-liquid phase separation for active zone formation and plasma membrane attachment (Emperador-Melero et al., 2021; McDonald et al., 2020; Wu et al., 2019, 2021).

Decades of gene knockout and related studies have uncovered loss-of-function phenotypes for individual active zone proteins. In essence, these studies established that each protein, in one way or another, participates in the control of each key exocytotic parameter. For example, roles in vesicle docking and priming have been described for RIM (Calakos et al., 2004; Deng et al., 2011; Han et al., 2011; Kaeser et al., 2011; Koushika et al., 2001), Munc13 (Aravamudan et al., 1999; Augustin et al., 1999; Deng et al., 2011; Imig et al., 2014; Richmond et al., 1999), Liprin-α (Emperador-Melero et al., 2021; Spangler et al., 2013; Wong et al., 2018), ELKS (Dong et al., 2018; Held et al., 2016; Kawabe et al., 2017; Matkovic et al., 2013), Piccolo/Bassoon (Parthier et al., 2018), and RIM-BP (Brockmann et al., 2019). Similarly, the control of Ca^2+^ secretion-coupling is mediated by the same proteins, with established roles for RIM, RIM-BP, Bassoon, and ELKS (Acuna et al., 2015; Davydova et al., 2014; Dong et al., 2018; Grauel et al., 2016; Han et al., 2011; Kaeser et al., 2011; Kittel et al., 2006; Liu et al., 2014, 2011). A true mechanistic understanding of the active zone, however, has been difficult to achieve. This is in part because reconstitution assays, powerful for untangling mechanisms of the fusion reaction itself (Hu et al., 2003; Ma et al., 2013; Tucker et al., 2004), are not possible for the active zone due to its molecular complexity. Furthermore, the redundancy of the scaffolding and release mechanisms has made it challenging to distinguish effects on active zone assembly and function, and the steps of build-up of release machinery for exocytosis of a vesicle have remained uncertain. Ultimately, it has not been possible to define which of the many candidate mechanisms at the active zone suffice to drive fast, action potential-triggered release.

Simultaneous conditional knockout of RIM and ELKS leads to disassembly of the active zone with loss of RIM, ELKS, RIM-BP, Piccolo, Bassoon and Munc13, a near-complete absence of vesicle docking, and a strong reduction in action-potential evoked exocytosis (Wang et al., 2016). Remarkably, general features of synaptic structure including the formation of boutons, accumulation of vesicles, and generation of synaptic contacts remain intact. This has established that RIM and ELKS form a scaffolding complex that holds the active zone together.

Here, we use this active zone disruption through RIM+ELKS knockout to develop an approach for reconstructing hallmark functions of these secretory machines within synapses. Our overall goal was to develop a deep mechanistic understanding of active zone function and to define which of the many mechanisms are sufficient to drive release. We experimentally identify the positioning of activated Munc13 to vesicles as a mechanism that induces fusion-competence. Notably, vesicles can be rendered release-competent by Munc13 without tethering them to the target membrane. Docking of these vesicles next to Ca^2+^ channels was required, however, to restore action potential-triggering of release. We achieved this using a single artificial fusion-protein consisting of the RIM zinc finger domain and Ca_V_β4-subunits, which leads to recovery of fusion strength, speed and spatial precision after active zone disruption. These findings establish the minimal requirements for active zone function, define key protein domains sufficient to mediate these requirements, and identify mechanisms that drive assembly of this minimal release machinery. Our work provides proof-of-concept for how an 80 kDa protein can be used to bypass the need for a megadalton-sized protein machine.

## Results

### RIM restores synaptic structure and function after active zone disruption

We first used stimulated emission depletion (STED) superresolution microscopy to evaluate active zone disruption induced by conditional knockout of RIM and ELKS, which strongly impaired extent, speed and precision of neurotransmitter release (Wang et al., 2016). Cultured hippocampal neurons with floxed alleles for RIM1, RIM2, ELKS1 and ELKS2 were infected with cre-expressing lentiviruses to generate knockout (cKO^R+E^) neurons or with control viruses (to generate control^R+E^ neurons). We then used a previously established workflow (Held et al., 2020; de Jong et al., 2018; Nyitrai et al., 2020; Wong et al., 2018) to assess active zone structure at 15-19 days in vitro (DIV). We identified side-view synapses in immunostainings by a bar-like postsynaptic density (PSD, marked by PSD-95, STED) that was aligned at one edge of a synaptic vesicle cloud (Synaptophysin, confocal), and assessed localization of target proteins (STED) relative to these markers in line profiles (Figs. S1A-S1D). The cKO^R+E^ synapses had disrupted active zones with near-complete loss of ELKS2, RIM1 and Munc13-1, and strong reductions in Bassoon and RIM-BP2 (Figs. 1A-1K). In addition, we observed a partial loss of Ca_V_2.1 (Figs. 1L, 1M), the α1-subunit of the voltage-gated Ca^2+^ channels that trigger release at these synapses (Held et al., 2020). Removal of these active zone complexes, however, did not affect the PSDs (marked by PSD-95, Fig. 1C), and led to increased levels of Liprin-α3 (Figs. S1E, S1F), a protein that connects active zone assembly pathways to structural plasticity (Emperador-Melero et al., 2021; Wong et al., 2018).

**Figure 1.**
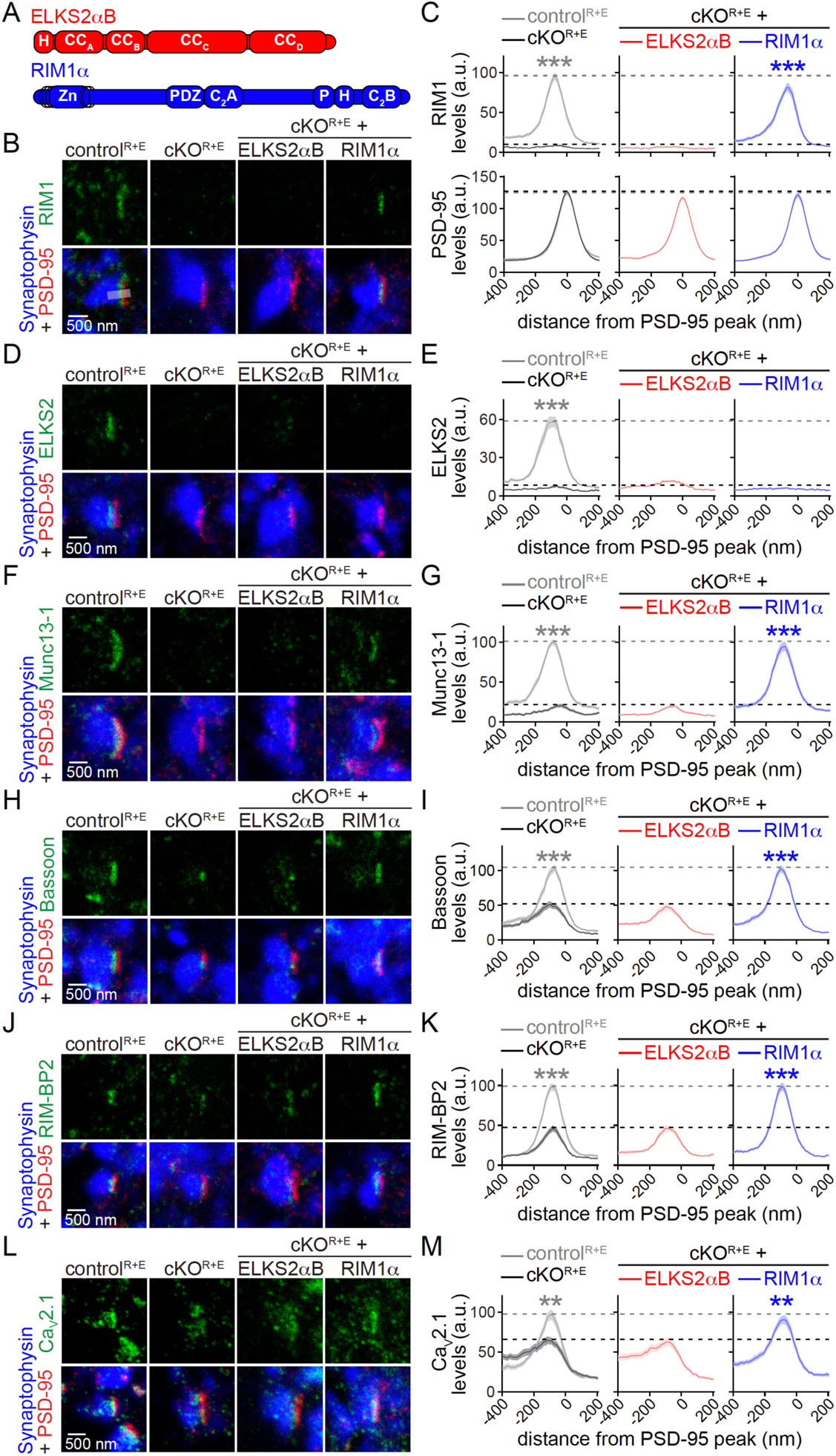
RIM restores active zone structure after knockout of RIM and ELKS. (**A**) Schematic of HA-tagged ELKS2αB and RIM1α that were expressed in cKO^R+E^ neurons using lentiviruses. H: HA tag, CC: coiled-coil region, Zn: zinc finger domain, PDZ: PDZ domain, C_2_A and C_2_B: C_2_ domains, P: proline rich (PxxP-) motif. (**B, C**) Sample STED images (B) and quantification (C) of side-view synapses in cultured hippocampal neurons after conditional knockout of RIM and ELKS (cKO^R+E^), stained for RIM1 (STED), PSD-95 (STED), and Synaptophysin (confocal) in control^R+E^ and cKO^R+E^ synapses, and in cKO^R+E^ synapses after lentiviral re-expression of ELKS2αB or RIM1α. Peak position and levels (C) were analyzed in line profiles (600 nm x 200 nm, grey shaded area in B) positioned perpendicular to the center of PSD-95 ‘bars’ and aligned to the PSD-95 peak, dotted lines mark control^R+E^ (grey) and cKO^R+E^ (black) levels for comparison, n = 60 synapses/3 independent cultures per condition. (**D-M**) Same as (B, C), but stained for ELKS2 (D, E), Munc13-1 (F, G), Bassoon (H, I), RIM-BP2 (J, K) or Ca_V_2.1 (L, M), n = 60/3 per condition. PSD-95 levels were assessed for all experiments as in C and outcomes were similar, but are not shown for simplicity. Data are mean ± SEM; \*\**P* < 0.01, \*\*\**P* < 0.001 compared to cKO^R+E^ as determined by two-way ANOVA followed by Dunnett’s multiple comparisons post-hoc tests. For STED analyses workflow, assessment of Liprin-α3, and assessment of rescue protein expression by Western blotting, see Fig. S1; for independent confocal microscopic experiments, see Fig. S2.

With the overall goal to rebuild active zone function using the minimally required protein domains and interactions, we first tested whether either RIM or ELKS mediate recovery of active zone structure and function on their own. We re-expressed RIM1α or ELKS2αB using lentiviruses (Figs. 1A, S1G), and found that RIM1α was targeted correctly (Figs. 1B, 1C) and was able to reestablish normal levels and active zone positioning of Munc13-1, Bassoon, RIM-BP2, Ca_V_2.1 and Liprin-α3 (Figs. 1F-1M, S1E, S1F). In contrast, rescue ELKS2αB did not localize to the target membrane area (Figs. 1D, 1E) and did not restore other proteins (Figs. 1F-1M, S1E, S1F), even though it was expressed efficiently (Fig. S1G). To assess protein levels upon rescue with an independent approach and to determine whether RIM-mediated active zone protein recruitment depends on the levels of RIM expression, we performed additional confocal microscopy experiments. Notably, re-expression of low or high RIM1α levels mediated recovery of the other proteins dose-dependently. Higher levels of RIM1α at synapses were driving the recruitment of higher levels of Munc13-1, Ca_V_2.1, and RIM-BP2 (Fig. S2), indicating that RIM is able to titrate the presynaptic levels of interacting active zone proteins.

**Figure 2.**
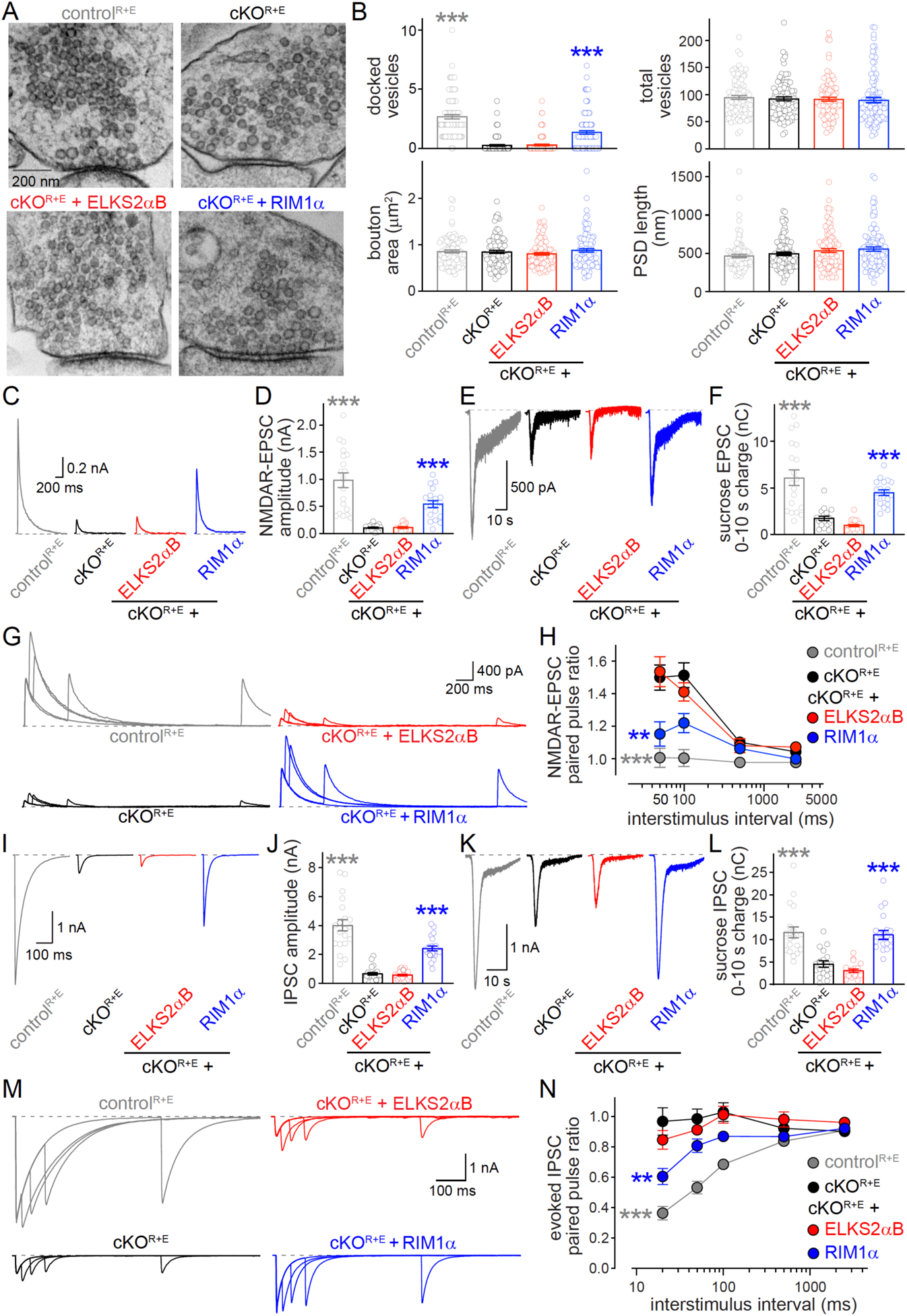
RIM restores active zone functions after active zone disruption. (**A, B**) Sample images (A) and analyses (B) of synapses of high-pressure frozen neurons analyzed by electron microscopy. Quantification was performed on single sections, control^R+E^ n = 96 synapses/2 independent cultures, cKO^R+E^ n = 100/2, cKO^R+E^ + ELKS2αB n = 100/2, and cKO^R+E^ + RIM1α n = 98/2. (**C, D**) Sample traces (C) and quantification (D) of EPSCs evoked by focal electrical stimulation, control^R+E^ n = 19 cells/3 independent cultures, cKO^R+E^ n = 18/3, cKO^R+E^ + ELKS2αB, n = 18/3, cKO^R+E^ + RIM1α n = 19/3. (**E, F**) Sample traces (E) and quantification (F) of EPSCs triggered by hypertonic sucrose, the first 10 s of the EPSC were quantified to estimate the RRP, n = 19/3 per condition. (**G, H**) Sample traces (G) and quantification (H) of EPSC paired pulse ratios to estimate p, n = 16/3 per condition. (**I, J**) Sample traces (I) and quantification (J) of IPSCs evoked by focal electrical stimulation, control^R+E^ n = 20/3, cKO^R+E^ n = 21/3, cKO^R+E^ + ELKS2αB n = 21/3, and cKO^R+E^ + RIM1α n = 22/3. (**K**, **L**) Sample traces (K) and quantification (L) of IPSCs triggered by hypertonic sucrose, the first 10 s of the IPSC were quantified to estimate the RRP, n = 19/3 per condition. (**M**, **N**) Sample traces (M) and quantification (N) of IPSC paired pulse ratios, n = 15/3 per condition. Data are mean ± SEM; \*\**P* < 0.01, \*\*\**P* < 0.001 compared to cKO^R+E^ as determined by Brown-Forsythe ANOVA followed by Games-Howell’s multiple comparisons post-hoc test (B, docked vesicles), Brown-Forsythe ANOVA followed by Dunnett’s T3 multiple comparisons post-hoc test (D), Kruskal-Wallis followed by Dunn’s multiple comparisons post-hoc tests (F, J and L), or by two-way ANOVA followed by Dunnett’s multiple comparisons post-hoc tests (H and N). For analyses of EPSCs in low extracellular Ca^2+^ or with competitive NMDAR antagonists, see Fig. S3.

We next tested whether RIM1α re-expression restored key active zone functions, synaptic vesicle docking and release (Fig. 2). To assess vesicle docking and synaptic ultrastructure, we fixed neurons with high pressure-freezing and analyzed electron microscopic images of synapses. Most cKO^R+E^ synapses lacked docked vesicles entirely (assessed as vesicles for which the electron density of the vesicle membrane merges with that of the active zone target membrane), but other parameters including PSD length, bouton size and total vesicle numbers were unaffected. RIM1α restored vesicle docking to 51% of its initial levels, while ELKS2αB expression did not improve docking (Figs. 2A, 2B).

The active zone controls synaptic strength by generating a readily releasable pool (RRP) of vesicles and by setting the release probability p of each RRP vesicle. We measured synaptic strength and estimated these constituents, p and RRP, at both excitatory and inhibitory synapses using electrophysiology (Figs. 2C-2N). p is inversely proportional to the ratio of release in response to paired pulses at short interstimulus intervals (Zucker and Regehr, 2002), and application of hypertonic sucrose was used to estimate the RRP (Kaeser and Regehr, 2017; Rosenmund and Stevens, 1996). RIM1α mostly restored excitatory (Figs. 2C-2H) and inhibitory (Figs. 2I-2N) synaptic strength, and both RRP and p were recovered to a large extent at both synapse types. In contrast, ELKS2αB had no rescue activity on its own, consistent with the STED and electron microscopy data. Excitatory evoked transmission was monitored via NMDA receptors (NMDARs) to avoid confounding effects of network activity triggered by AMPA receptor activation. Decreasing initial p by lowering extracellular Ca^2+^ or the use of low affinity NMDAR antagonists confirmed that paired pulse ratios provide an accurate estimate of changes in p as a consequence of genetic manipulations under our conditions (Fig. S3). Together, these data establish that RIM is an important presynaptic organizer for the control of active zone protein levels, positioning and function.

**Figure 3.**
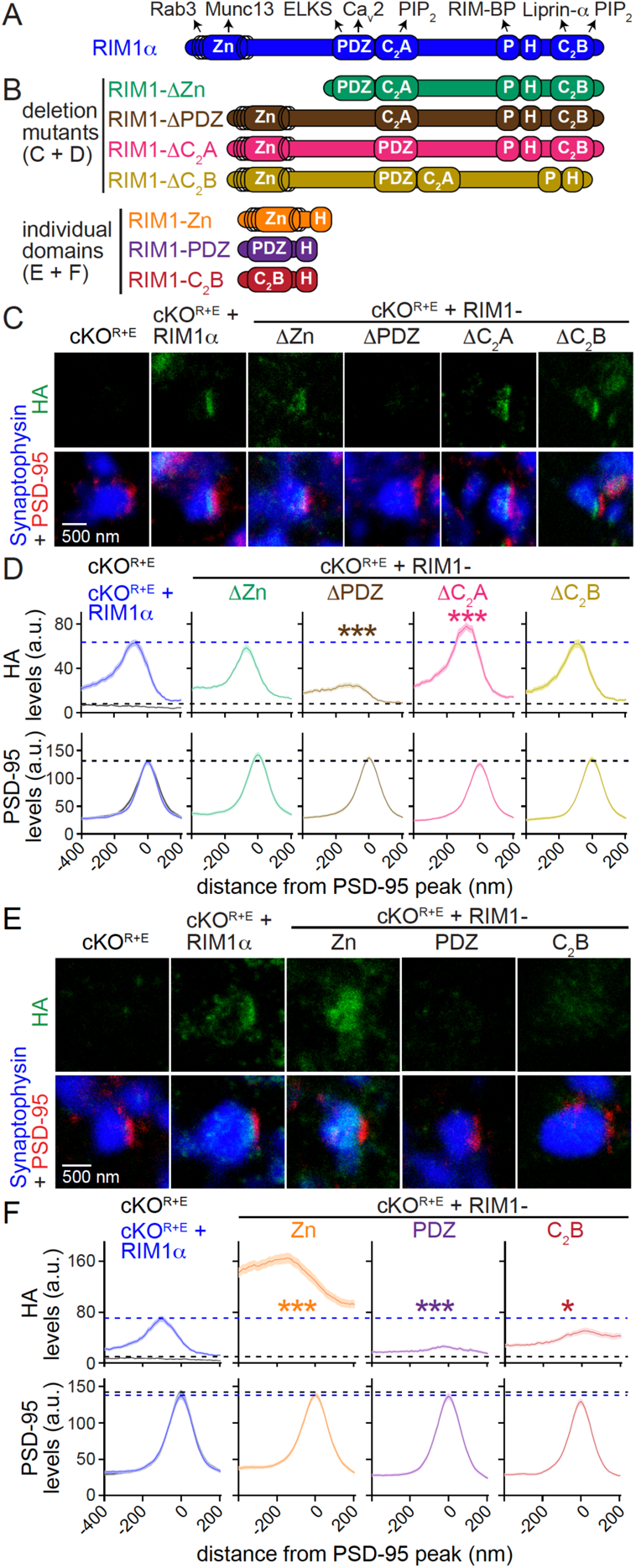
RIM PDZ domains mediate RIM active zone localization while zinc fingers associate with synaptic vesicles. (**A, B**) Schematic of key presynaptic interactors of full-length RIM1α (A) and of RIM1 deletion mutants or individual domains (B) that were expressed in cKO^R+E^ neurons. (**C, D**) Sample STED images (C) and quantification (D) of side-view synapses stained for HA (STED), PSD-95 (STED), and Synaptophysin (confocal), dotted lines mark cKO^R+E^ (black) and cKO^R+E^ + RIM1α (blue) levels for comparison, n = 60 synapses/3 independent cultures per condition. (**E, F**) Same as (C, D), but for individual domains, n = 60/3 per condition. Data are mean ± SEM; \**P* < 0.05, \*\*\**P* < 0.001 compared to cKO^R+E^ + RIM1α as determined by two-way ANOVA followed by Dunnett’s tests. For assessment of rescue expression by Western blotting, see Figs. S4A and S4B.

### RIM zinc fingers localize to the synaptic vesicle cloud

For building a minimal recovery system, we next needed to distinguish between RIM domains that mediate active zone targeting of RIM from those that are important for its functions in scaffolding other proteins and in mediating vesicle docking and release. We generated lentiviral constructs (Figs. 3A, 3B) in which we either deleted individual RIM domains (RIM1-ΔZn, -ΔPDZ, -ΔC_2_A and -ΔC_2_B) or that contained only one domain at a time (RIM1-Zn, -PDZ and -C_2_B; the tested C_2_A domain constructs were not efficiently expressed and hence C_2_A domains could not be assessed in these experiments). cKO^R+E^ neurons were transduced with each individual virus for rescue, and each protein was efficiently expressed (Figs. S4A, S4B).

Assessment of RIM active zone targeting using STED line profile analyses revealed that the PDZ domain was necessary for RIM target membrane localization after active zone disruption, as removing the PDZ domain abolished RIM active zone targeting (Figs. 3C, 3D). Other domain deletions did not impair RIM localization. Notably, no single domain of RIM was targeted to the plasma membrane opposed to the PSD when expressed alone (Figs. 3E, 3F). Hence, while multiple RIM domains cooperated for RIM active zone targeting, the RIM PDZ domain is essential for such targeting. It is noteworthy that in neurons lacking only RIM rather than RIM and ELKS, most active zone proteins remain localized to synapses, and the PDZ domain is not essential for RIM localization to nerve terminals (de Jong et al., 2018; Kaeser et al., 2011). In active zone disrupted cKO^R+E^ synapses, this redundancy is lost and the PDZ domain is essential (Figs. 3C, 3D). Hence, the highly interconnected active zone protein networks rely on redundant scaffolding mechanisms that include ELKS (Held and Kaeser, 2018).

Notably, the RIM1 zinc finger alone, while not localized to the active zone, was strongly enriched within nerve terminals (Figs. 3E, 3F). RIM1-Zn localization highly overlapped with the synaptic vesicle protein Synaptophysin, suggesting that this domain may be associated with vesicles when expressed on its own. Since the RIM zinc finger interacts with the vesicular GTPases Rab3 and Rab27 (Dulubova et al., 2005; Fukuda, 2003; Wang et al., 1997), it is likely that RIM1-Zn associates through these interactions with synaptic vesicles. The complementary protein fragment, a version of RIM that lacks the zinc finger domain termed RIM1-ΔZn, localized to the active zone area apposed to the PSDs (Figs. 3C, 3D), establishing that the zinc finger domain is not required for the synaptic delivery of RIM.

### RIM1 zinc fingers recruit Munc13 for establishing release-competence of non-docked vesicles

The differential localization of RIM1-Zn (to vesicles) and RIM1-ΔZn (to the target membrane) may be related to their roles in release. Previous studies in RIM knockout synapses have suggested that RIM zinc finger domains prime synaptic vesicles while the C-terminal domains within RIM1-ΔZn tether Ca_V_2 channels and interact with the target membrane for fast release triggering (Deng et al., 2011; de Jong et al., 2018; Kaeser et al., 2011). We tested these models by assessing the molecular roles (recruitment of Munc13 and Ca_V_2s) and functional roles (priming, docking and releasing of vesicles) of RIM1-Zn and RIM1-ΔZn after active zone disruption.

Strikingly, RIM1-Zn co-recruited Munc13 in a pattern mimicking the wide-spread localization of RIM1-Zn (Figs. 4B, 4C). In contrast, RIM1-ΔZn, which contains the RIM scaffolding domains (PDZ, C_2_A, PxxP, C_2_B) and localizes to the active zone area (Figs. 3A-3D), was unable to enhance Ca_V_2s or Munc13 (Figs. 4B-4E). This suggests that RIM1-Zn, which binds to Munc13 and Rab3 (Betz et al., 2001; Dulubova et al., 2005), recruits Munc13 to synapses, stabilizes it, and turns is into a protein associated with synaptic vesicles rather than the target membrane. We next analyzed high pressure-frozen neurons using electron microscopy from the same rescue conditions. Both RIM1-Zn and RIM1-ΔZn lacked rescue activity, and synaptic vesicles remained undocked in both conditions (Figs. 4F, 4G). In conclusion, Munc13, a protein implicated in synaptic vesicle docking (Imig et al., 2014; Siksou et al., 2009), was not targeted to the presynaptic plasma membrane upon RIM1-Zn re-expression but was instead associated with the vesicle cloud. Recovering its presence in the nerve terminal was not sufficient to mediate docking.

**Figure 4.**
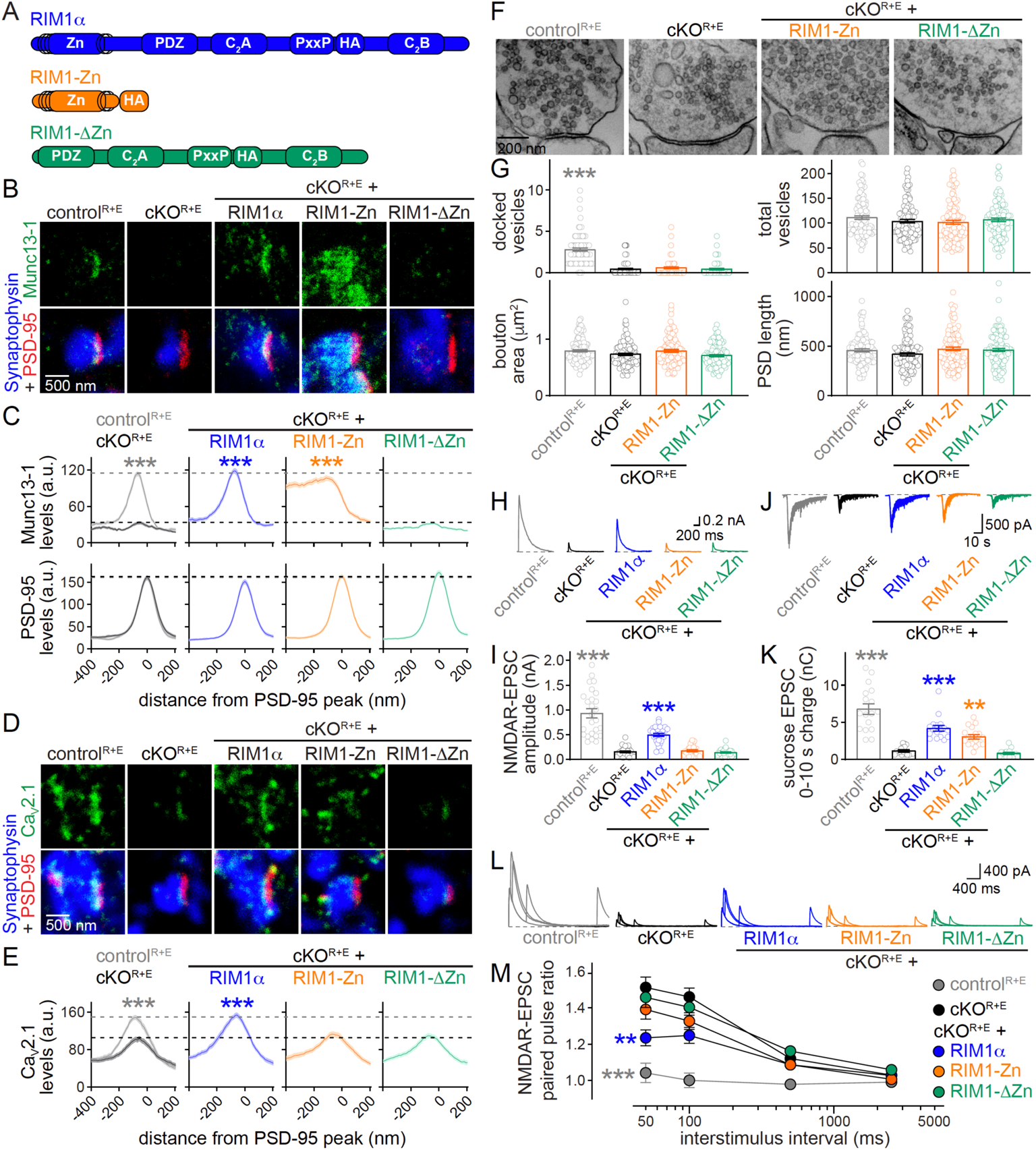
RIM zinc fingers render undocked vesicles release-competent through Munc13 recruitment. (**A**) Schematic of rescue proteins. (**B, C**) Sample STED images (B) and quantification (C) of side-view synapses stained for Munc13-1 (STED), PSD-95 (STED), and Synaptophysin (confocal), dotted lines mark control^R+E^ (grey) and cKO^R+E^ (black) levels for comparison, n = 60 synapses/3 independent cultures per condition. (**D, E**) Same as (B, C), but for Ca_V_2.1, n = 60/3 per condition. PSD-95 levels were assessed as in C and outcomes were similar, but are not shown for simplicity. (**F, G**) Sample electron microscopic images (F) and analyses (G) of synapses of high-pressure frozen neurons, control^R+E^ n = 105 synapses/2 independent cultures, cKO^R+E^ n = 110/2, cKO^R+E^ + RIM1-Zn n = 105/2, and cKO^R+E^ + RIM1-ΔZn n = 102/2. It was not possible to include RIM1α full-length rescue in all electron microscopic experiments due to the laborious nature of processing and analyses. As essential controls, we included control ^R+E^ and cKO^R+E^ neurons in each electron microscopic experiment. (**H, I**) Sample traces (H) and quantification (I) of EPSCs evoked by focal electrical stimulation, n = 26 cells/4 independent cultures per condition. (**J, K**) Sample traces (J) and quantification (K) of EPSCs triggered by hypertonic sucrose to estimate the RRP, n = 17/3 per condition. (**L, M**) Sample traces (L) and quantification (M) of paired pulse ratios to estimate p, n = 25/4 per condition. Data are mean ± SEM; \*\**P* < 0.01, \*\*\**P* < 0.001 compared to cKO^R+E^ as determined by two-way ANOVA followed by Dunnett’s tests (C, E and M), Brown-Forsythe ANOVA followed by Games-Howell’s multiple comparisons post-hoc test (G, docked vesicles), or by Kruskal-Wallis analysis followed by Dunn’s tests (I and K). For recordings of IPSCs, see Figs. S4C-S4H.

Electrophysiological recordings of excitatory (Figs. 4H-4M) and inhibitory (Figs. S4C-S4H) transmission revealed that RIM1-Zn and RIM1-ΔZn failed to restore action potential-triggered synaptic transmission and release probability of excitatory synapses (Figs. 4H-4M), and only mild rescue of these parameters was observed at inhibitory synapses (Figs. S4C-S4H). This is different from rescue experiments after RIM knockout only (instead of cKO^R+E^), where RIM1-ΔZn is sufficient to restore Ca^2+^ entry and mediates an increase in p (Deng et al., 2011; Kaeser et al., 2011). Hence, RIM C-terminal scaffolding domains need ELKS or N-terminal RIM sequences (Figs. 1 and 2) to execute their roles in release, but are sufficient to mediate target membrane localization of RIM (Figs. 3C, 3D).

Remarkably, however, RIM1-Zn strongly enhanced vesicle fusogenicity measured via application of hypertonic sucrose, nearly as efficiently as full-length RIM1α (Figs. 4J, 4K, S4E, S4F). These data support the model that RIM zinc fingers activate Munc13 for vesicle priming by recruiting and monomerizing it via binding to Munc13 C_2_A domains (Camacho et al., 2017; Deng et al., 2011) and – strikingly – this function can be executed on non-docked vesicles distant from the target membrane. These vesicles, however, are inaccessible to action potential-triggering. We conclude that undocked vesicles can become release-competent by positioning activated Munc13 on them. Hence, Munc13 enhances fusogenicity even if it is not localized to release sites at the target membrane, and these “molecularly” primed vesicles do not need to be docked. This may explain why some priming remains when the active zone is disrupted and docking is abolished (Wang et al., 2016), as some Munc13 may be recruited to vesicles via direct interactions (Quade et al., 2019).

### Docking of release-competent vesicles to Ca^2+^ channels restores fast release in the absence of active zone scaffolds

With the goal to selectively rebuild active zone mechanisms without restoring the vast scaffolding structure, we aimed at positioning the release-competent vesicles close to Ca^2+^ entry. We screened eight fusion-proteins of the RIM zinc finger domain to other proteins or protein fragments associated with the target membrane (Fig. S5A). Fusions with Ca_V_β Ca^2+^ channel subunits appeared to efficiently restore evoked transmission and release probability (Figs. S5B-S5E), suggesting that they may do so by co-localizing vesicle priming and Ca^2+^-entry. We selected the fusion of the RIM1 zinc finger to Ca_V_β4 (Fig. 5A, β4-Zn) for a full characterization because of its strong tendency to rescue and because endogenous Ca_V_β4 is localized to active zones (Figs. S5F, S5G). β4-Zn was efficiently expressed (Fig. S5H) and concentrated in a bar-shaped structure at the target membrane (Figs. 5A-5C). The β4-Zn protein efficiently recruited Munc13-1 to the target membrane (Figs. 5D, 5E), and also enhanced Ca_V_2.1 active zone levels back to control levels (Figs. 5F, 5G). The effects on Ca_V_2.1 were notably absent when Ca_V_β4 or RIM1-Zn were expressed on their own. Importantly, β4-Zn did not enhance or restore the levels of the other active zone scaffolds as assessed by staining and quantification of Bassoon and RIM-BP2 (Figs. 5H-5K). Hence, β4-Zn targets the priming-complex of the RIM zinc finger and Munc13 to Ca^2+^ channels in the absence of the megadalton-sized scaffolding network that consists of full-length RIM, ELKS, RIM-BP and Bassoon.

**Figure 5.**
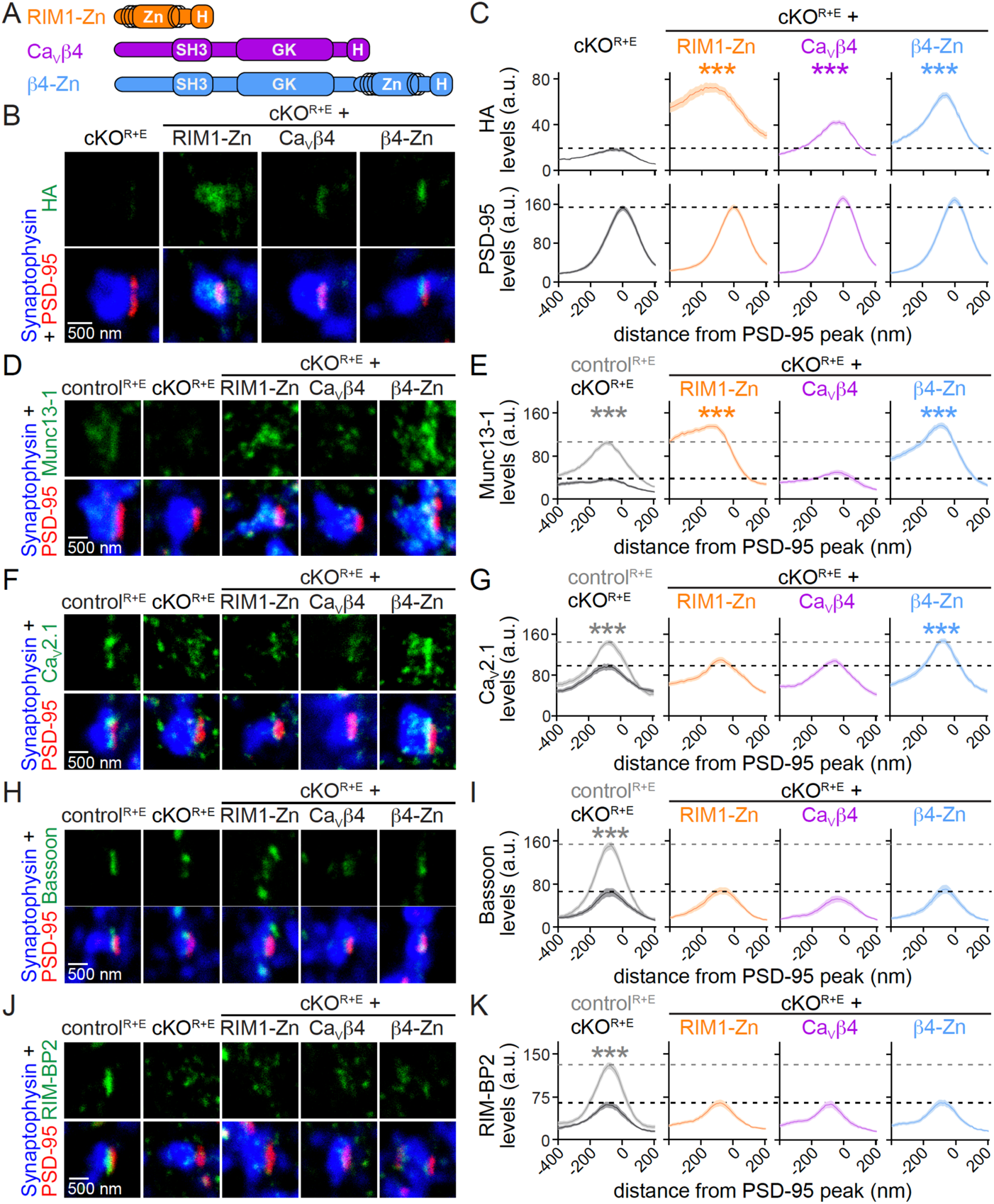
A Ca_V_β4-RIM1 zinc finger fusion-protein recruits priming machinery close to Ca^2+^ channels. (**A**) Schematic of rescue proteins. (**B, C**) Sample STED images (B) and quantification (C) of side-view synapses stained for HA to visualize rescue proteins (STED), PSD-95 (STED), and Synaptophysin (confocal), dotted lines mark cKO^R+E^ (black) levels for comparison, n = 60 synapses/3 independent cultures per condition. (**D-K**) Same as (B, C), but stained for Munc13-1 (D, E), Ca_V_2.1 (F, G), Bassoon (H, I), and RIM-BP2 (J, K), n = 60/3 in each condition. PSD-95 levels were assessed for all experiments as in C and outcomes were similar, but are not shown for simplicity, dotted lines mark control^R+E^ (grey) and cKO^R+E^ (black) levels for comparison. Data are mean ± SEM; \*\*\**P* < 0.001 compared to cKO^R+E^ as determined by two-way ANOVA followed by Dunnett’s tests. For assessment of various rescue fusion-proteins, STED analyses of Cavβ4, and expression levels of rescue proteins by Western blot, see Fig. S5.

When we assessed these synapses using electron microscopy, we found that β4-Zn fully restored synaptic vesicle docking (Figs. 6A, 6B). This is different from expression of Ca_V_β4 or RIM1-Zn alone, which did not enhance docking (Figs. 6A, 6B), and much more robust than rescue efficacy of full-length RIM1α (Figs. 2A, 2B). In electrophysiological recordings, we detected a full recovery of excitatory (Figs. 6C, 6D) and inhibitory (Figs. 6I, 6J) synaptic transmission. This included restoration of RRP (Figs. 6E, 6F, 6K, 6L) and p (Figs. 6G, 6H, 6M, 6N) back to levels indistinguishable from control^R+E^ neurons at both types of synapses. Hence, β4-Zn expression results in Munc13 recruitment and vesicle docking to the active zone close to Ca^2+^ channels such that extent and spatiotemporal precision of release are fully restored.

**Figure 6.**
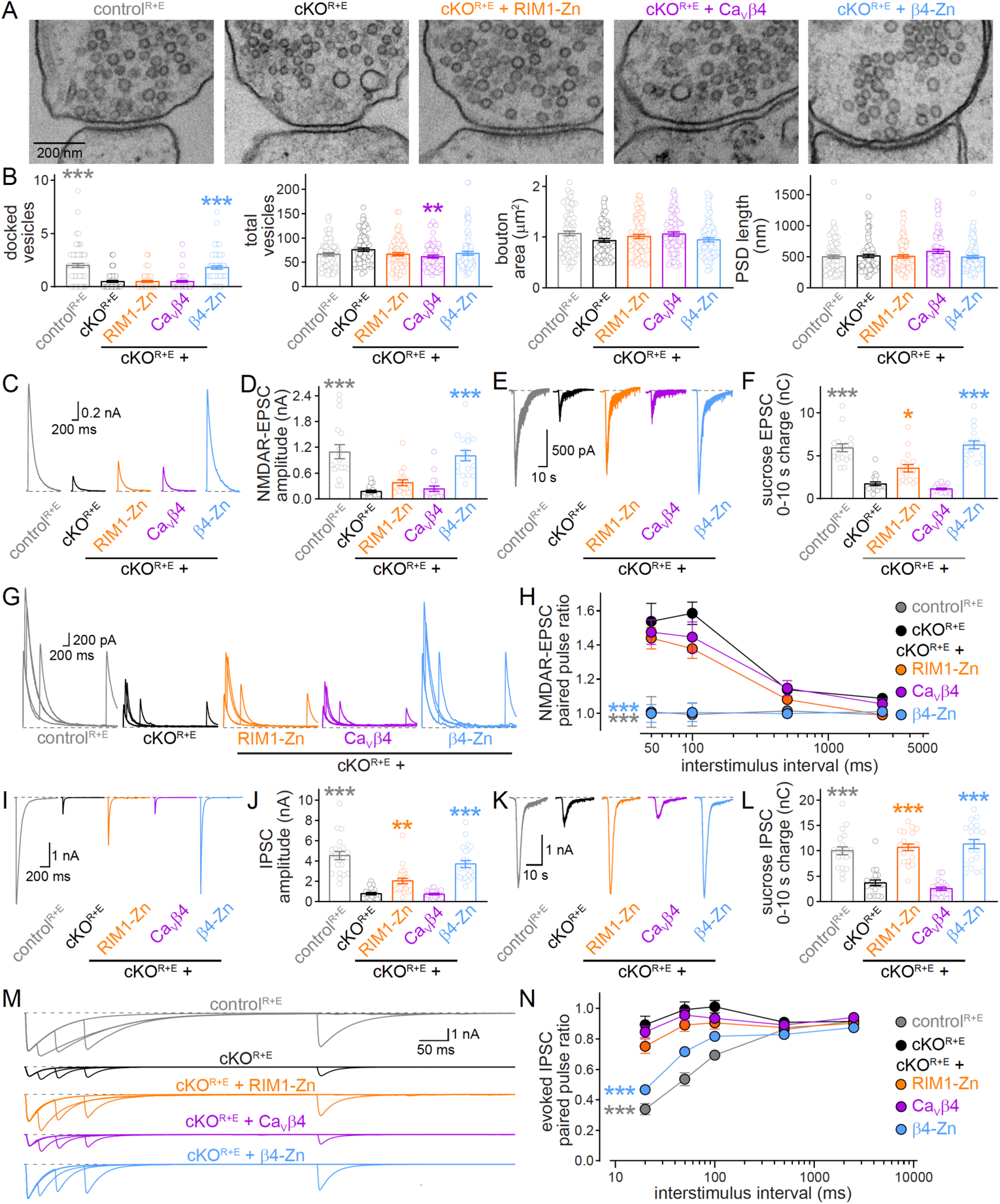
β4-Zn reconstitutes vesicle docking and release. (**A, B**) Sample electron microscopic images (A) and analyses (B) of synapses of high-pressure frozen neurons, control^R+E^ n = 83 synapses/2 independent cultures, cKO^R+E^ n = 85/2, cKO^R+E^ + RIM1-Zn n = 84/2, cKO^R+E^ + Ca_V_β4 n = 83/2 and cKO^R+E^ + β4-Zn n = 87/2. (**C, D**) Sample traces (C) and quantification (D) of EPSCs evoked by focal electrical stimulation, control^R+E^ n = 17 cells/3 independent cultures, cKO^R+E^ n = 17/3, cKO^R+E^ + RIM1-Zn n = 18/3, cKO^R+E^ + Ca_V_β4 n = 18/3 and cKO^R+E^ + β4-Zn n = 17/3. (**E, F**) Sample traces (E) and quantification (F) of the EPSC triggered by hypertonic sucrose, the first 10 s of the EPSC were quantified to estimate the RRP, n = 17/3 per condition. (**G, H**) Sample traces (G) and quantification (H) of EPSC paired pulse ratios to estimate p, control^R+E^ n = 16/3, cKO^R+E^ n = 16/3, cKO^R+E^ + RIM1-Zn n = 17/3, and cKO^R+E^ + Ca_V_β4 n = 16/3, cKO^R+E^ + β4-Zn n = 17/3. (**I, J**) Sample traces (I) and quantification (J) of IPSCs evoked by focal electrical stimulation, control^R+E^ n = 21/4, cKO^R+E^ n = 21/4, cKO^R+E^ + RIM1-Zn n = 24/4, cKO^R+E^ + Ca_V_β4 n = 22/4, and cKO^R+E^ + β4-Zn n = 22/4. (**K**, **L**) Sample traces (K) and quantification (L) of IPSC triggered by hypertonic sucrose, the first 10 s of the IPSC were quantified to estimate the RRP, n = 20/3 per condition. (**M**, **N**) Sample traces (M) and quantification (N) of IPSC paired pulse ratios, control^R+E^ n = 20/4, cKO^R+E^ n = 19/4, cKO^R+E^ + RIM1-Zn n = 20/4, cKO^R+E^ + Ca_V_β4 n = 19/4, and cKO^R+E^ + β4-Zn n = 20/4. Data are mean ± SEM; \**P* < 0.05, \*\**P* < 0.01, \*\*\**P* < 0.001 compared to cKO^R+E^ as determined by Brown-Forsythe ANOVA followed by Games-Howell’s multiple comparisons post-hoc test (B, docked vesicles), one-way ANOVA followed by Dunnett’s multiple comparisons post-hoc tests (B, total vesicles), Kruskal-Wallis followed by Dunn’s tests (D, F, J and L) or by two-way ANOVA followed by Dunnett’s tests (H and N).

Importantly, Ca_V_β4 or RIM1-Zn alone did not mediate these functions except for partial RRP recovery by RIM1-Zn (Fig. 4, S4, 6), and β4-Zn recovered docking and release without reinstating broader active zone scaffolds (Fig. 5). These data establish that vesicle docking close to Ca^2+^ channels enhances p of vesicles that are primed by RIM1-Zn and Munc13. They further suggest that vesicle docking close to Ca_V_2s might not only enhance p, but also stabilize Ca^2+^ channels. Alternatively, interactions of Ca_V_β4 with vesicles and Munc13 may help Ca_V_2 delivery to synapses.

To test the overall model that β4-Zn restores synaptic strength through docking of release-competent vesicles close to Ca^2+^ channels, we introduced K144E+K146E point mutations into the RIM1 zinc finger domain of β4-Zn (generating β4-Zn^K144/6E^). It was previously established that this mutation selectively abolishes binding of the RIM zinc finger to Munc13 (Deng et al., 2011; Dulubova et al., 2005). β4-Zn^K144/6E^ was efficiently expressed and localized to the active zone area of the plasma membrane (Figs. 7A-7C, S6A). Abolishing Munc13 binding in β4-Zn^K144/6E^ resulted in a loss of Munc13-1 recruitment to the target membrane (Figs. 7D, 7E).

**Figure 7.**
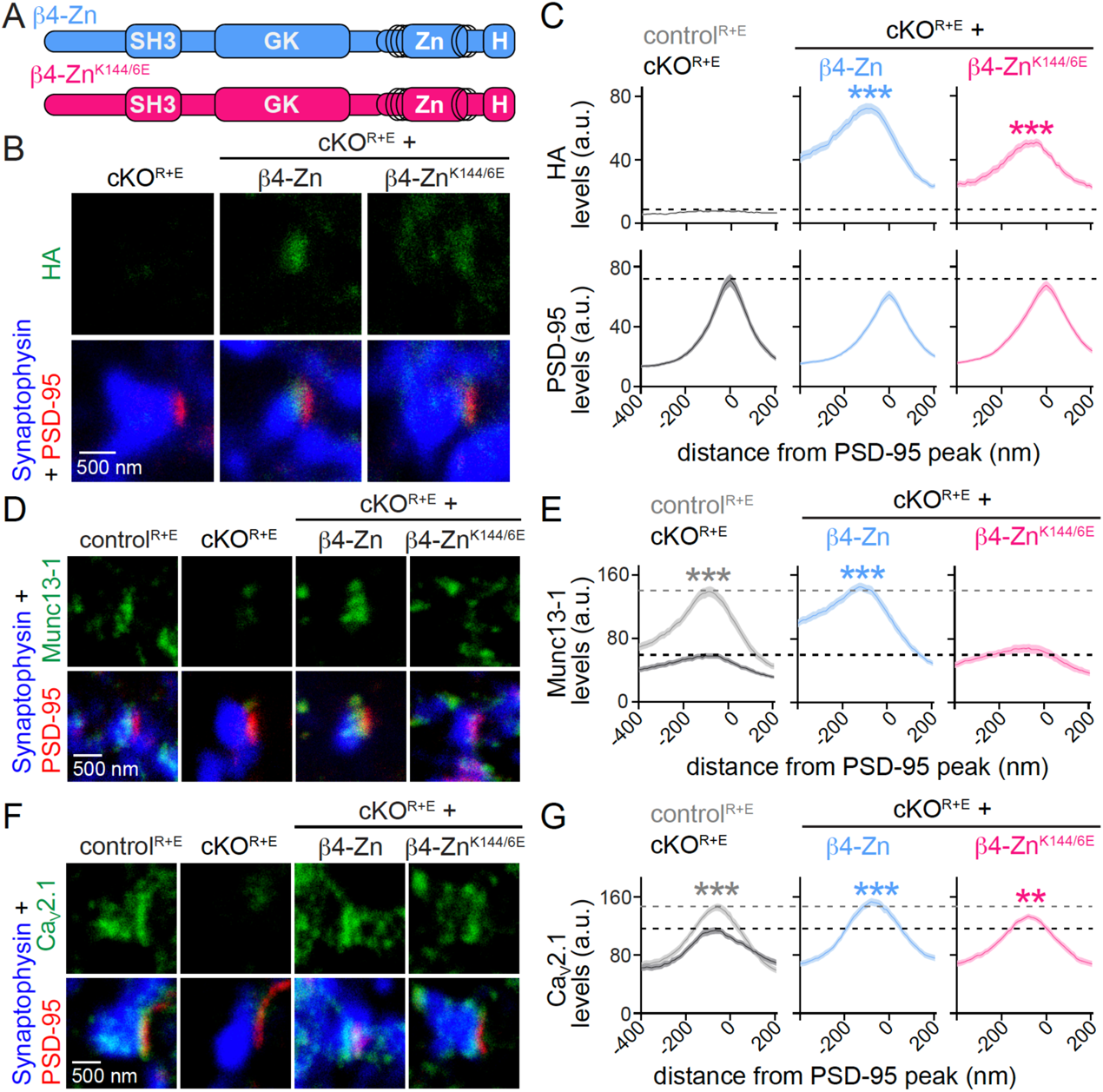
Binding of β4-Zn to Munc13 mediates presynaptic Munc13 recruitment. (**A**) Schematic of rescue proteins. (**B, C**) Sample STED images (C) and quantification (D) of side-view synapses stained for HA (STED), PSD-95 (STED), and Synaptophysin (confocal) in cKO^R+E^ synapses, and in cKO^R+E^ synapses after re-expression of β4-Zn or β4-Zn^K144/6E^, dotted lines mark cKO^R+E^ levels for comparison, n = 60 synapses/3 independent cultures per condition. (**D-G**) Same as (B, C), but stained for Munc13-1 (D, E) and Ca_V_2.1 (F, G), n = 60/3 in each condition. PSD-95 levels were assessed for all experiments as in C, but are not shown for simplicity, dotted lines mark control^R+E^ (grey) and cKO^R+E^ (black) levels for comparison. Data are mean ± SEM; \*\**P* < 0.01, \*\*\**P* < 0.001 compared to cKO^R+E^ as determined by two-way ANOVA followed by Dunnett’s tests. For assessment of rescue protein expression by Western blotting, see Fig. S6A.

While β4-Zn^K144/6E^ was still sufficient to mediate some enhancement of Ca_V_2.1 levels, it appeared less efficient than in β4-Zn (Figs. 7F, 7G), quantitatively matching with the somewhat lower β4-Zn^K144/6E^ active zone levels (Figs. 7B, 7C). Notably, disrupting binding of β4-Zn to Munc13 completely abolished rescue of vesicle docking (Figs. 8A, 8B) and of synaptic strength, RRP and p at both excitatory (Figs. 8C-8H) and inhibitory (Figs. S6B-S6G) synapses. These data strongly support the model that β4-Zn restores exocytosis via recruitment of Munc13 and vesicle docking. Altogether, our results establish that the active zone protein network can be bypassed with an 80 kDa protein that docks fusion-competent vesicles close to Ca^2+^ channels, and the vast active zone scaffolding mechanisms are not necessary for fusion itself.

**Figure 8.**
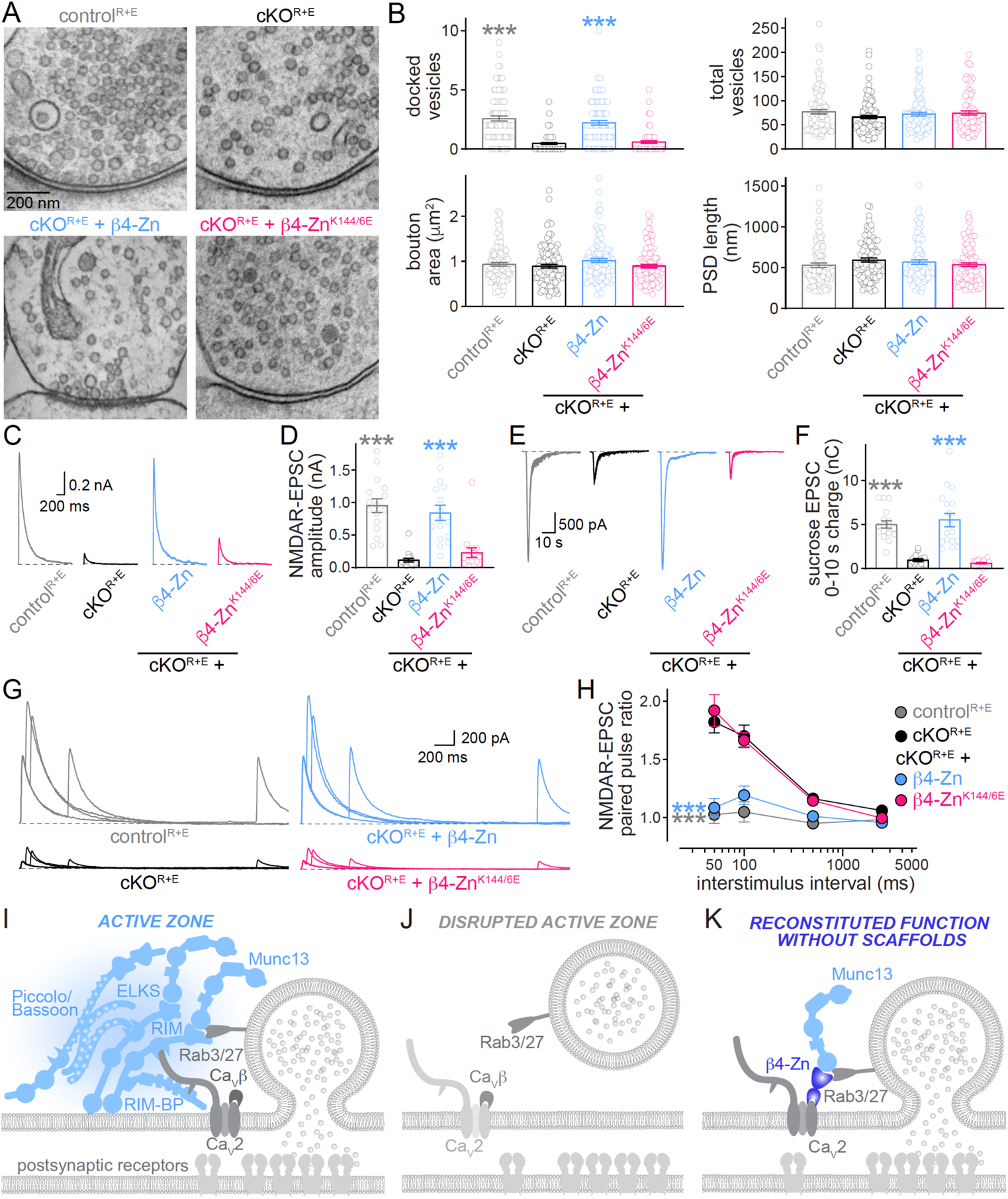
Binding of β4-Zn to Munc13 is essential for restoring vesicle docking and release. (**A, B**) Sample electron microscopic images (A) and analyses (B) of synapses of high-pressure frozen neurons, control^R+E^ n = 100 synapses/2 independent cultures, cKO^R+E^ n = 99/2, cKO^R+E^ + β4-Zn n = 99/2 and cKO^R+E^ + β4-Zn^K144/6E^ n = 99/2. (**C, D**) Sample traces (C) and quantification (D) of EPSCs evoked by focal electrical stimulation, n = 16 cells/4 independent cultures per condition. (**E, F**) Sample traces (E) and quantification (F) of the EPSC triggered by hypertonic sucrose, the first 10 s of the EPSC were quantified to estimate the RRP, n = 17/3 per condition. (**G, H**) Sample traces (G) and quantification (H) of paired pulse ratios to estimate p, n = 16/4 per condition. (**I-K**) Schematic shows the active zone, a protein network assembled through liquid-liquid phase separation, which controls extent, speed and spatial precision of synaptic vesicle release (I). Knockout of RIM and ELKS leads to loss of active zone scaffolds and of vesicle docking, and impaired release (J). Reconstitution of synaptic function can be achieved in the absence of the active zone scaffolding network. Step-wise reconstitution of active zone function was accomplished by reintroduction of the RIM zinc finger, which localizes to synaptic vesicles, co-recruits endogenous Munc13, and renders non-docked vesicles highly fusogenic (Figures 3 and 4). When fused onto the Ca^2+^ channel subunit Ca_V_β4 (K, Figures 5–7), vesicle docking and rapid exocytosis are restored. This reconstitution approach bypasses the need for the active zone scaffolding network. Data are mean ± SEM; \*\*\**P* < 0.001 compared to cKO^R+E^ as determined by Brown-Forsythe ANOVA followed by Games-Howell’s multiple comparisons post-hoc test (B, docked vesicles), Kruskal-Wallis followed by Dunn’s tests (D and F) or by two-way ANOVA followed by Dunnett’s tests (H). For IPSC β4-Zn^K144/6E^ rescue data, see Figs. S6B-S6G.

## Discussion

The active zone is a molecular machine that is important for synaptic signaling (Emperador-Melero and Kaeser, 2020; Südhof, 2012), and many brain disorders are associated with mutations in active zone proteins or defective active zone function (Benarroch, 2013; Bucan et al., 2009; Johnson et al., 2003; Krumm et al., 2015; O’Roak et al., 2012; Thevenon et al., 2013; Verhage and Sørensen, 2020). However, understanding its mechanisms and restoring its functions has remained challenging because of its molecular complexity. Mouse knockout experiments have revealed that each active zone protein contributes to each active zone function, and it has remained uncertain which proteins and which specific mechanisms drive its roles in release. Here, we develop a reconstitution approach within a synapse after we remove the active zone protein machinery. Our work establishes that synaptic vesicle release can be restored by positioning the RIM zinc finger, a single, small protein domain that recruits the priming protein Munc13, to the Ca^2+^ channels that mediate release triggering. Remarkably, doing so bypasses the need for the complex active zone scaffolding network. Our work further reveals that if the RIM zinc finger is localized to vesicles, Munc13 is recruited and can render these vesicles fusion-competent in the absence of docking. Ultimately, these findings present a straightforward way to restore efficacy and spatiotemporal precision of neurotransmitter release at central synapses with a single, small protein.

### Flexible order of priming and docking as a vesicle is prepared for release

Our results mechanistically define the two fundamental presynaptic processes: vesicle fusogenicity can be generated by activated Munc13 independent of its active zone positioning, and Ca^2+^-secretion coupling is mediated by docking of Munc13-associated synaptic vesicles next to Ca^2+^ channels. Past studies have discovered that these processes rely on many proteins, and each active zone protein has contributed to each active zone function (Acuna et al., 2015; Aravamudan et al., 1999; Augustin et al., 1999; Brockmann et al., 2019; Davydova et al., 2014; Deng et al., 2011; Dong et al., 2018; Emperador-Melero et al., 2021; Grauel et al., 2016; Held et al., 2016; Imig et al., 2014; Kaeser et al., 2011; Kawabe et al., 2017; Kittel et al., 2006; Koushika et al., 2001; Lipstein et al., 2013; Liu et al., 2014, 2011; Matkovic et al., 2013; Richmond et al., 1999; Schoch et al., 2002; Varoqueaux et al., 2002; Wong et al., 2018; Zhen and Jin, 1999). These findings reflect that the active zone is a complex protein network with built-in redundancy, and knockout studies may lead to alterations of the entire network and not necessarily reveal highly specific mechanisms of isolated proteins. Hence, it has been difficult to define which proteins and mechanisms drive vesicle priming, docking and release. Our approach establishes that these functions can be executed in the absence of most of these proteins. Vesicle priming can be almost entirely mediated by the RIM1 zinc finger domain, which is sufficient to recruit, stabilize and activate Munc13. When this mechanism is positioned close to Ca^2+^ channels, release is restored.

Models of neurotransmitter release propose that vesicle docking either precedes vesicle priming or occurs simultaneously with it (Hammarlund et al., 2007; Imig et al., 2014; Rosenmund and Stevens, 1996; Schikorski and Stevens, 2001; Sudhof, 2004), and that Munc13 mediates both roles through the control of SNARE complex assembly (Basu et al., 2005; Imig et al., 2014; Ma et al., 2013; Siksou et al., 2009; Südhof, 2012). In view of this literature, it is surprising that vesicle fusogenicity can be generated in the absence of docking (Fig. 4), dissociating the linear docking-priming model that relies on SNARE-complex assembly. Our data instead reveal that the generation of fusion-competent vesicles and vesicle docking are molecularly separable processes, and that vesicles away from the active zone can be activated for fusion. We propose that the rate limiting step is not SNARE-complex assembly, as this necessitates the coincidence of docking and priming, but instead the availability and activation of Munc13. This is supported by the observation that fusion-competent vesicles can be generated by positioning Munc13 on undocked vesicles (Figs. 3 and 4), by the notion that some fusion-competent vesicles remain when Munc13 is displaced from the active zone (Wang et al., 2016), and by the finding that knockout of RIM, which is upstream of Munc13’s role in vesicle priming, leads to strong impairments in RRP (Calakos et al., 2004; Deng et al., 2011; Han et al., 2011). Hence, Munc13 mediates both synaptic vesicle docking and priming (Augustin et al., 1999; Imig et al., 2014; Varoqueaux et al., 2002), but generating fusion competence and vesicle docking are molecularly separable. We propose that, while many RRP vesicles are docked (Borges-Merjane et al., 2020; Imig et al., 2014; Schikorski and Stevens, 2001), the presence and activation of Munc13 embody the bottleneck for vesicle priming. Munc13-mediated SNARE complex-assembly may not be rate-limiting for vesicle release and can occur before or during fusion.

### Streamlined active zone assembly bypasses the need for complex scaffolds

Mechanisms and hierarchy of active zone protein recruitment have remained difficult to establish. Our work reveals that the RIM PDZ domain is important for recruitment of RIM to the active zone, as removing it prevents RIM active zone targeting. Furthermore, our data indicate that RIM drives recruitment of presynaptic protein machinery. RIM re-expression restores levels of all other active zone proteins and of Ca^2+^ channels, and, remarkably, does so in a dose-dependent manner such that more RIM drives the presence of more Munc13, Ca^2+^ channels and other active zone proteins. Recent work proposed that liquid-liquid phase separation of RIM, RIM-BP and Ca_V_2s mediates active zone assembly (Wu et al., 2019, 2021). While our work does not directly test this model, it is consistent with it, but indicates that phase condensation of RIM and RIM-BP into liquid droplets may not be necessary for neurotransmitter release, because simply fusing the RIM zinc finger domain to Ca_V_β4 restores release in the absence of most RIM and RIM-BP sequences necessary for phase condensation. In principle it is possible that other liquid phases, which may or may not incorporate Ca_V_2s, could be at play. In this context, it is interesting that Liprin-α3 levels at the active zone increase upon active zone disruption. Liprin-α proteins undergo phase condensation, and participate in the regulation of active zone structure (Emperador-Melero et al., 2021; McDonald et al., 2020). It is possible, and perhaps likely, that the two phases compete or are in equilibrium with one another at a synapse, and that removing one enhances the other. This is supported by the enhanced presence of Liprin-α3 upon disruption of the active zone protein complex between RIM, ELKS, RIM-BP, Ca_V_2s, Munc13 and Bassoon (Fig. 1). Conversely, at Liprin-α2/3 knockout synapses, enhanced levels of Ca_V_2 proteins are present (Emperador-Melero et al., 2021). RIM may link the two phases together or participate in both, as its active zone recruitment is decreased upon Liprin-α ablation.

An interesting observation is that the artificial β4-Zn fusion protein enhances Ca_V_2 active zone levels together with restoring vesicle docking and release. One possibility is that vesicle docking stabilizes the Ca_V_2 protein complex at active zones. This model is supported by the observation that abolishing the docking function of the β4-Zn fusion protein by preventing its binding to Munc13 appears to revert this function at least partially. Another possibility is that the β4-Zn fusion protein enhances the delivery of Ca_V_2s to the active zone, and that Munc13 binding is required for this function. Ultimately, our data may suggest that a stable release site contains a docked vesicle, that release sites that do not contain docked vesicles are subject to dynamic rearrangements, and that Ca_V_2s of unoccupied release sites may be more mobile (Schneider et al., 2015). An alternative model is that Ca_V_2s and exocytotic protein machinery such as Munc13 are in different proteins complexes (Rebola et al., 2019). In this model RIM proteins would participate in distinct assemblies: one may define secretory sites and contain at least RIM and Munc13 (Deng et al., 2011; Emperador-Melero et al., 2021; Reddy-Alla et al., 2017; Sakamoto et al., 2018; Tang et al., 2016), and another controls Ca^2+^ channel clustering and contains RIM, RIM-BP and Ca_V_2s (Acuna et al., 2016; Held et al., 2020; Hibino et al., 2002; Kaeser et al., 2011; Kushibiki et al., 2019; Liu et al., 2011; Oh et al., 2021). In this model, proteins like RIM or ELKS could bridge the complexes, and our reconstitution would account for both functions (Fig. 8K). Future studies should address these models.

In aggregate, our data suggest that the release machinery assembly requirements are remarkably simple: RIM zinc fingers recruit Munc13 to prime vesicles, and if positioned next to Ca_V_2 channels, these vesicles can be rapidly and precisely released (Figs. 8I-8K). We propose that synaptic strength can be controlled through reconstituting these key mechanisms, and that the other protein domains and liquid phases mediate regulatory functions (Emperador-Melero and Kaeser, 2020; Emperador-Melero et al., 2021). Some secretory systems, for example those for striatal dopamine release, may make use of these relatively simple, streamlined mechanisms (Banerjee et al., 2021; Liu et al., 2018).

### A small protein for rebuilding the function of a complex machine

Neurotransmitter secretion is often impaired in brain disease, ranging from highly specific associations of gene mutations in active zone proteins to more generalized breakdown of secretion and transmitter signaling (Benarroch, 2013; Bucan et al., 2009; Johnson et al., 2003; Krumm et al., 2015; O’Roak et al., 2012; Thevenon et al., 2013; Verhage and Sørensen, 2020). Advances in AAV-based gene therapy strategies have spurred new hope for developing treatments for brain disorders (Hudry and Vandenberghe, 2019; Sun and Roy, 2021). However, a key limitation is that synaptic and secretory genes often exceed the packaging size of AAVs (Wu et al., 2010). A recent way to work past this limitation is the use of dual or triple AAVs for expression of fragments that are then spliced to generate whole proteins, for example for restoration of hearing (Akil et al., 2019; Al-Moyed et al., 2019). Another possibility is to find smaller proteins to restore function. Our reconstitution approach identifies a single 80 kDa-protein, the Ca_V_β4-RIM zinc finger fusion, that is well within packaging limits of gene therapy viruses (Hudry and Vandenberghe, 2019; Wu et al., 2010). It is remarkable that this relatively small protein can strongly enhance synaptic efficacy and is sufficient to mediate spatiotemporal precision of release. Our approach may serve as proof-of-concept for reconstructing functions of a complex molecular machine with relatively simple “pieces”. Ultimately, this finding may be leveraged to develop new approaches for enhancing transmitter secretion in neurological and endocrine diseases.

## Acknowledgments

We thank J. Wang and C. Qiao for technical support, Drs. E. Raviola, W. Regehr, R. Held and H. Nyitrai for insightful discussions and help with setting up experimental approaches, all members of the Kaeser laboratory for feedback, and Drs. M. Verhage and J. Broeke for a MATLAB macro to analyze electron microscopic images. This work was supported by grants from the NIH (R01NS083898 and R01MH113349 to PSK), the Armenise Harvard Foundation (to PSK), and an NSF graduate research fellowship (DGE1144152 to S.S.H.W.). We acknowledge the Neurobiology Imaging Facility (supported by a P30 Core Center Grant P30NS072030), and the Electron Microscopy Facility at Harvard Medical School.

## Author contributions

Conceptualization, CT, SSHW, and PSK; Methodology, CT, SSHW, and GdN; Formal Analysis, CT, SSHW, GdN and PSK; Investigation, CT, SSHW and GdN; Resources, CT, SSHW; Writing-Original Draft, CT and PSK; Writing-Review & Editing, CT, GdN and PSK; Supervision, PSK; Funding Acquisition, PSK.

## Materials and Methods

### Mice

The quadruple homozygote floxed mice for RIM1αβ (Kaeser et al., 2008) (RRID: IMSR_JAX:015832), RIM2αβγ (Kaeser et al., 2011) (RRID: IMSR_JAX:015833), ELKS1α (Liu et al., 2014) (RRID: IMSR_JAX:015830) and ELKS2α (Kaeser et al., 2009) (RRID: IMSR_JAX:015831) were previously described (Wang et al., 2016). All animal experiments were performed according to institutional guidelines at Harvard University.

### Cell culture and lentiviral infection

Primary mouse hippocampal cultures were generated from newborn conditional quadruple floxed pups as described before (Held et al., 2020; Wang et al., 2016). Mice were anesthetized on ice slurry within 24 h after birth and the hippocampus was dissected out. Cells were dissociated and plated onto glass coverslips in tissue culture medium composed of Mimimum Essential Medium (MEM) with 0.5% glucose, 0.02% NaHCO_3_, 0.1 mg/mL transferrin, 10% Fetal Select bovine serum (Atlas Biologicals FS-0500-AD), 2 mM L-glutamine, and 25 µg/mL insulin. Cultures were maintained in a 37 °C-tissue culture incubator, and after ~24 h the plating medium was exchanged with growth medium composed of MEM with 0.5% glucose, 0.02% NaHCO_3_, 0.1 mg/mL transferrin, 5% Fetal Select bovine serum (Atlas Biologicals FS-0500-AD), 2% B-27 supplement (Thermo Fisher 17504044), and 0.5 mM L-glutamine. At DIV3 or DIV4, depending on growth, 50% or 75% of the medium were exchanged with growth medium supplemented with 4 µM Cytosine β-D-arabinofuranoside (AraC) to inhibit glial cell growth. Lentiviruses expressing EGFP-tagged cre recombinase (to generate cKO^R+E^ neurons, made using pFSW EGFP cre) or a truncated, enzymatically inactive EGFP-tagged cre protein (to generate control^R+E^ neurons, made using pFSW EGFP Δcre) were produced in HEK293T cells by Ca^2+^-phosphate transfection. Expression in lentiviral constructs was driven by the human Synapsin promoter to restrict expression to neurons (Liu et al., 2014; Wang et al., 2016) except for the RIM1α_high_ condition (which was done using FUGW-RIM1α with a ubiquitin promoter). For cre-expressing and control virus, neurons were infected with HEK cell supernatant at DIV5 as described (Liu et al., 2014; Wang et al., 2016). For rescue with the various protein variants (ELKS1αB, ELKS2αB, RIM1α, RIM1 mutants, Ca_V_β4, β4-Zn and other RIM1-Zn fusion constructs), neurons were infected with rescue virus at DIV3 (a virus made using pFSW without a cDNA inserted in the multiple cloning site was used in the control conditions instead of a rescue virus) and with cre or Δcre virus at DIV5. Analyses were performed at DIV15-19.

### Rescue constructs

For full-length RIM1α (all residue numbering is provided according to Uniprot ID Q9JIR4), the open reading frame (ORF) was subcloned into lentiviral backbones and expression was driven by either a synapsin promoter (pFSW RIM1α-HA, p592) for lower expression or a ubiquitin promoter (pFUGW RIM1α-HA, p591, described in (de Jong et al., 2018)) for higher expression. The synapsin promoter was used in all other rescue constructs.

For all experiments, RIM zinc finger refers to residues M1-D213, RIM PDZ to H597-R705, RIM C_2_A to Q754-Q882, and RIM C_2_B to G1447-S1615. All RIM1 individual domains (pFSW RIM1-Zn-HA, p654; pFSW RIM1-PDZ-HA, p648; pFSW RIM1-C2B-HA, p647) and domain deletion mutants (pFSW RIM1-ΔZn-HA, p640; pFSW RIM1-ΔPDZ-HA, p639; pFSW RIM1-ΔC2A-HA, p637; pFSW RIM1-ΔC2B-HA, p638) span or lack these residues, except for the pFSW RIM1-ΔZn-HA, which spans H597-S1615. In RIM1α and in domain deletion mutants, an HA-tag was inserted between residues E1379-S1380. In RIM1 individual domains, an HA-tag was inserted at the C-terminus. The splice variant of full-length RIM1α was lacking alternatively spliced exons (N83-W105, H1084-R1169, A1207-T1378) identical to previous experiments (Deng et al., 2011; de Jong et al., 2018; Kaeser et al., 2011; Tang et al., 2016). For pFSW HA-ELKS1αB (p311) and pFSW HA-ELKS2αB (p314), an HA-tag was inserted at the N-terminus (Held et al., 2016; Nyitrai et al., 2020). The plasmids for expression of zinc finger fusion-proteins were newly generated based on the following cDNAs: pMT2 Ca_V_β1b GFP (gift from Annette Dolphin obtained through Addgene, plasmid # 89893; http://addgene.org/89893; RRID:Addgene_89893 (Page et al., 2016)), Cavβ3 (gift from Diane Lipscombe; http://addgene.org/26574 RRID:Addgene_26574) and pMT2 Ca_V_β4 (gift from Annette Dolphin obtained through Addgene, plasmid # 107426; http://addgene.org/107426; RRID:Addgene_107426 (Page et al., 2016)). The cDNAs of Liprin-α3 (Wong et al., 2018), Ca_V_2.1 (Held et al., 2020) and RIM1-Zn^K144/6E^ (Deng et al., 2011) were described before. For pFSW β4-Zn-HA (p661), an HA-tag followed by RIM1-Zn was inserted at the C-terminus of Ca_V_β4, with the stop codon in Ca_V_β4 and start codon in RIM1-Zn deleted. For all other RIM1-Zn fusion-proteins, similar strategies were used as shown in Fig. S5A.

### STED imaging

Neurons cultured on 0.17 mm thick 12 mm diameter (#1.5) coverslips were washed two times with warm PBS, and then fixed in 4% PFA for 10 min unless noted otherwise. For Ca_V_2.1 staining, cultures were fixed in 2% PFA + 4% sucrose (in PBS) for 10 min. After fixation, coverslips were rinsed twice in PBS + 50 mM glycine, then permeabilized in PBS + 0.1% Triton X-100 + 3% BSA (TBP) for 1 hour. Primary antibodies were diluted in TBP and stained for 24-48 h at 4 °C. The following primary antibodies were used: guinea pig anti-Synaptophysin (1:500, RRID: AB_1210382, A106), mouse anti-PSD-95 (1:200, RRID: AB_10698024, A149), rabbit anti-RIM1 (1:500, RRID: AB_887774, A58), rabbit anti-ELKS2α (serum E3-1029, 1:100, custom made, A136, (Held et al., 2016)), rabbit anti-Munc13-1 (1:500, RRID: AB_887733, A72), rabbit anti-Ca_V_2.1 (1:200, RRID: AB_2619841, A46), rabbit anti-RIM-BP2 (1:500, RRID: AB_2619739, A126), rabbit anti-Liprin-α3 (serum 4396, 1:2000, gift from Dr. T. Südhof, A35), rabbit anti-Synaptophysin (1:500, RRID: AB_887905, A64), guinea pig anti-Bassoon^C^ (C-terminal, 1:500, RRID: AB_2290619, A67) and mouse anti-HA (1:500, RRID: AB_2565006, A12). After primary antibody staining, coverslips were rinsed twice and washed 3-4 times for 5 minutes in TBP. Alexa Fluor 488 (anti-guinea pig, RRID: AB_2534117, S3; anti-rabbit, RRID: AB_2576217, S5; anti-mouse IgG1, RRID: AB_2535764, S7), 555 (anti-mouse IgG2a, RRID: AB_1500824, S20), and 633 (anti-rabbit, RRID: AB_2535731, S33; anti-guinea pig, RRID: AB_2535757, S34) conjugated antibodies were used as secondary antibodies at 1:200 (Alexa Fluor 488 and 555) or 1:500 (Alexa Fluor 633) dilution in TBP, incubated for 24 h at 4 °C followed by rinsing two times and washing 3-4 times 5 minutes in TBP. Stained coverslips were post-fixed for 10 minutes with 4% PFA in PBS (for Ca_V_2.1 staining, 4% PFA + 4% sucrose in PBS was used for post-fixation), rinsed two times in PBS + 50 mM glycine and once in deionized water, and air-dried and mounted on glass slides. STED images were acquired with a Leica SP8 Confocal/STED 3X microscope with an oil immersion 100x 1.44 numerical aperture objective and gated detectors as described in (Wong et al., 2018). 46.51 x 46.51 µm^2^ areas were taken as regions of interest (ROIs) and were scanned at a pixel density of 4096 x 4096 (11.358 nm/pixel). Alexa Fluor 633, Alexa Fluor 555, and Alexa Fluor 488 were excited with 633 nm, 555 nm and 488 nm using a white light laser at 2-5% of 1.5 mW laser power. The Alexa Fluor 633 channel was acquired first in confocal mode using 2x frame averaging. Subsequently, Alexa Fluor 555 and Alexa Fluor 488 channels were acquired in STED mode, depleted with 660 nm (50% of max power, 30% axial depletion) and 592 nm (80% of max power, 30% axial depletion) depletion lasers, respectively. Line accumulation (2-10x) and frame averaging (2x) were applied during STED scanning. Identical settings were applied to all samples within an experiment. Synapses within STED images were selected in side-view, defined as synapses that contained a synaptic vesicle cluster labeled with Synaptophysin and associated with an elongated PSD-95 bar along the edge of the vesicle cluster as described (Held et al., 2020; de Jong et al., 2018; Wong et al., 2018). For intensity profile analyses, side-view synapses were selected using only the PSD-95 signal and the vesicle signal for all experiments. An ROI was manually drawn around the PSD-95 signal and fit with an ellipse to determine the center position and orientation. An ~1200 nm long, 200 nm wide rectangle was then selected perpendicular and across the center of the elongated PSD-95 structure. Intensity profiles were obtained for all three channels within this ROI. To align individual profiles, the PSD-95 signal only was smoothened using a moving average of 5 pixels, and the smoothened signal was used to define the peak position of PSD-95. All three channels (vesicle marker, test protein, and smoothened PSD-95) were then aligned to the PSD-95 peak position and averaged across images. All analyses were performed on raw images without background subtraction, and all adjustments and were done identically for all experimental conditions. Representative images were brightness and contrast adjusted to facilitate inspection, and these adjustments were made identically for images within an experiment. The experimenter was blind to the condition/genotype for image acquisition and analyses.

### Confocal imaging of cultured neurons

Neurons cultured on glass coverslips were washed with warm PBS and fixed in PFA for 20 min. Neurons were the permeabilized in TBP for 1 h, and then incubated in primary antibodies at 4 °C overnight. The following primary antibodies were used: rabbit anti-RIM1 (1:1000, RRID: AB_887774, A58), rabbit anti-ELKSα (1:500, RRID: AB_869944, A55), rabbit anti-Munc13-1 (1:500, RRID: AB_887733, A72), rabbit anti-Ca_V_2.1 (1:1000, RRID: AB_2619841, A46), rabbit anti-RIM-BP2 (1:500, RRID: AB_2619739, A126), mouse anti-Bassoon (1:500, RRID: AB_11181058, A85), mouse anti-MAP2 (1:500, RRID: AB_477193, A108), rabbit anti-MAP2 (1:1000, RRID: AB_2138183, A139), guinea pig anti-Synaptophysin (1:500, RRID: AB_1210382, A106). After staining with primary antibodies, coverslips were rinsed twice and washed 3-4 times for 5 min in TBP. Alexa Fluor 488 (for detection of the protein of interest, anti-rabbit, RRID: AB_2576217, S5; anti-mouse IgG1, RRID: AB_2535764, S7), 546 (for detection of MAP2, anti-mouse IgG, RRID: AB_2535765, S12; anti-rabbit, RRID: AB_2534093, S16), and 633 (for detection of Synaptophysin, anti-guinea pig, RRID: AB_2535757, S34) conjugated secondary antibodies were used at 1:500 dilution in TBP. Secondary antibody staining was done for 2 h at room temperature followed by rinsing two times and washing 3-4 times 5 min in TBP. Coverslips were rinsed once with deionized water and mounted on glass slides. Images were taken on an Olympus FV1200 confocal microscope using identical settings per condition in a given experiment with a 60x oil-immersion objective and single confocal sections were analyzed in ImageJ. For quantitative analyses of synaptic protein levels, the Synaptophysin signal was used to define synaptic puncta as ROIs, and the signal intensity of the protein of interest was quantified within these ROIs. For each image, the “rolling ball” ImageJ plugin was set to a diameter of 1.4 µm for local background subtraction (Sternberg, 1983). Representative images were brightness and contrast adjusted to facilitate inspection, and adjustments were made identically across conditions. The experimenter was blind to the condition/genotype for image acquisition and analyses.

### Electrophysiology

Electrophysiological recordings in cultured hippocampal neurons were performed as described (Held et al., 2020; Liu et al., 2014; Wang et al., 2016) at DIV15-19. The extracellular solution contained (in mM): 140 NaCl, 5 KCl, 2 MgCl_2_, 1.5 CaCl_2_, 10 glucose, 10 HEPES-NaOH (pH 7.4, ~300 mOsm), for Figs. S3A-S3D, the CaCl_2_ concentration was 0.5 mM instead of 1.5 mM, all recordings were performed at room temperature (20 - 24 °C). To assess action potential-triggered excitatory transmission, NMDAR-mediated excitatory postsynaptic currents (EPSCs) were measured to avoid network activity induced by AMPA receptor activation. For NMDAR-EPSCs, picrotoxin (PTX, 50 µM) and 6-Cyano-7-nitroquinoxaline-2,3-dione (CNQX, 20 µM) were present in the extracellular solution, for Figs. S3E-S3H, 20 μM L- AP5 was added to the extracellular solution. Inhibitory postsynaptic currents (IPSCs) were recorded in the presence of D-amino-5-phosphonopentanoic acid (D-APV, 50 µM) and CNQX (20 µM) in the extracellular solution. Action potentials were elicited with a bipolar focal stimulation electrode fabricated from nichrome wire. Paired pulse ratios were calculated as the amplitude of the second PSC divided by the amplitude of the first PSC. The baseline value for the second PSC was taken immediately after the second stimulus artifact. For sucrose-induced EPSC recordings, TTX (1 µM), PTX (50 µM), and D-APV (50 µM) were added to the extracellular solution, and for sucrose-induced IPSC recordings, TTX (1 µM), CNQX (20 µM), and D-APV (50 µM) were added. The RRP was estimated by application of 500 mM sucrose in extracellular solution applied via a microinjector syringe pump for 10 s at a rate of 10 µl/min through a tip with an inner diameter of 250 µm. Glass pipettes were pulled at 2 - 5 MΩ and filled with intracellular solutions containing (in mM) for EPSC recordings: 120 Cs-methanesulfonate, 2 MgCl_2_, 10 EGTA, 4 Na_2_-ATP, 1 Na-GTP, 4 QX314-Cl, 10 HEPES-CsOH (pH 7.4, ~300 mOsm) and for IPSC recordings: 40 CsCl, 90 K-gluconate, 1.8 NaCl, 1.7 MgCl_2_, 3.5 KCl, 0.05 EGTA, 2 Mg-ATP, 0.4 Na_2_-GTP, 10 phosphocreatine, 4 QX314-Cl, 10 HEPES-CsOH (pH 7.2, ~300 mOsm). Cells were held at +40 mV for NMDAR-EPSC recordings and at −70 mV for sucrose EPSC, eIPSC and sucrose IPSC recordings. Access resistance was monitored during recordings and compensated to 3-5 MΩ, and cells were discarded if the uncompensated access exceeded 15 MΩ. Data were acquired at 5 kHz and lowpass filtered at 2 kHz with an Axon 700B Multiclamp amplifier and digitized with a Digidata 1440A digitizer. All data acquisition and analysis was done using pClamp10. For electrophysiological experiments, the experimenter was blind to the genotype throughout data acquisition and analysis.

### High-pressure freezing and electron microscopy

Neurons cultured on 6 mm matrigel-coated sapphire coverslips were frozen using a Leica EM ICE high-pressure freezer in extracellular solution containing (in mM): 140 NaCl, 5 KCl, 2 MgCl_2_, 2 CaCl_2_, 10 HEPES-NaOH (pH 7.4), 10 Glucose (~310 mOsm) with CNQX (20 µM), D-AP5 (50 µM) and PTX (50 µM) added to block synaptic transmission. After freezing, samples were first freeze-substituted (AFS2, Leica) in 1% glutaraldehyde, 1% osmium tetroxide, 1% water in anhydrous acetone as follows: −90 °C for 5 h, 5 °C per h to −20 °C, −20 °C for 12 h, and 10 °C per hour to 20 °C. Following freeze substitution, samples were Epon infiltrated, and baked for 48 h at 60 °C followed by 80 °C overnight before sectioning at 50 nm. For ultrathin sectioning, the sapphire coverslip was removed from the resin block by plunging the sample first in liquid nitrogen and followed by warm water several times until the sapphire was completely detached. The resin block containing the neurons was then divided into four pieces, and one piece was mounted for sectioning. Ultrathin sectioning was performed on a Leica EM UC7 ultramicrotome, and the 50 nm sections were collected on a nickel slot grid (2 x 1 mm) with a carbon coated formvar support film. The samples were counterstained by incubating the grids with 2% lead acetate solution for 10 seconds, followed by rinsing with distilled water. Images were taken with a transmission electron microscope (JEOL 1200 EX at 80 kV accelerating voltage) and processed with ImageJ. The total number of vesicles, the number of docked vesicles, the length of the PSD, and the area of the presynaptic bouton were analyzed in each section using a custom-written Matlab code. Bouton size was calculated from the measured perimeter of each synapse. Docked vesicles were defined as vesicles touching the presynaptic plasma membrane opposed to the PSD, and only vesicles for which the electron densities of the vesicular membrane and the presynaptic plasma membrane merged such that they were not separated by less electron dense space were considered docked. Due to the laborious nature of these experiments, it was not possible to include a RIM1α full-length rescue condition in each experiment. Instead, we always included control^R+E^ and cKO^R+E^ neurons as essential controls for comparison. Experiments and analyses were performed by an experimenter blind to the genotype.

### Western blotting

For assessment of rescue protein expression in cultured neurons, Western blotting was used to detect target proteins in cell lysates from select coverslips of every culture that was used for electrophysiology or electron microscopy. At DIV15-19, cultured neurons were harvested in 20 µl 1x SDS buffer per coverslip and run on standard SDS-Page gels followed by transfer on nitrocellulose membranes. Membranes were blocked in filtered 10% nonfat milk/5% goat serum for 1 h at room temperature and incubated with primary antibodies in 5% nonfat milk/2.5% goat serum overnight at 4 °C, and HRP-conjugated secondary antibodies (1:10,000, anti-mouse, RRID: AB_2334540; anti-rabbit, RRID: AB_2334589) were used. Anti-Synapsin or - β-actin antibodies were used as a loading controls. The following primary antibodies were used: rabbit anti-RIM1 (1:1000, RRID: AB_887774, A58), rabbit anti-ELKS2αB (1:500, RRID: AB_731499, A143), rabbit anti-Munc13-1 (1:1000, RRID: AB_887733, A72), rabbit anti-RIM1-Zn (1:500, gift from Dr T. Südhof, A148), mouse anti-HA (1:1000, RRID: AB_2565006, A12), mouse anti-Synapsin (1:4000, RRID: AB_2617071, A57), mouse anti-β-actin (1:2000, RRID: AB_476692, A127), mouse anti-Ca_V_β4 (1:50, RRID: AB_10671176, A123). For illustration in figures, images were adjusted for brightness and contrast to facilitate visual inspection.

### Statistics

Data are displayed as mean ± SEM, statistics were performed in GraphPad Prism 9, and significance is presented as * *P* < 0.05, ** *P* < 0.01, and *** *P* < 0.001. Parametric tests were used for normally distributed data (assessed by Shapiro-Wilk tests) or when sample size was n ≥ 30. One-way ANOVA followed by Dunnett’s multiple comparisons post-hoc tests were used for datasets with equal variance. When variances were unequal, Brown-Forsythe ANOVA followed by Games-Howell’s multiple comparisons post hoc tests (for n ≥ 50) or Dunnett’s T3 multiple comparisons post hoc tests (for n < 50) were used. For non-normally distributed data, nonparametric tests were used (Mann-Whitney tests or Kruskal-Wallis analysis of variance followed by Dunn’s multiple comparisons post-hoc tests). For paired pulse ratios, two-way ANOVA with Dunnett’s tests was used. For STED side-view analyses, two-way ANOVA with Dunnett’s tests was used on a 200 nm-window centered around the active zone peak. For each dataset, the specific tests used are stated in the figure legends.

---

**Figure S1.**
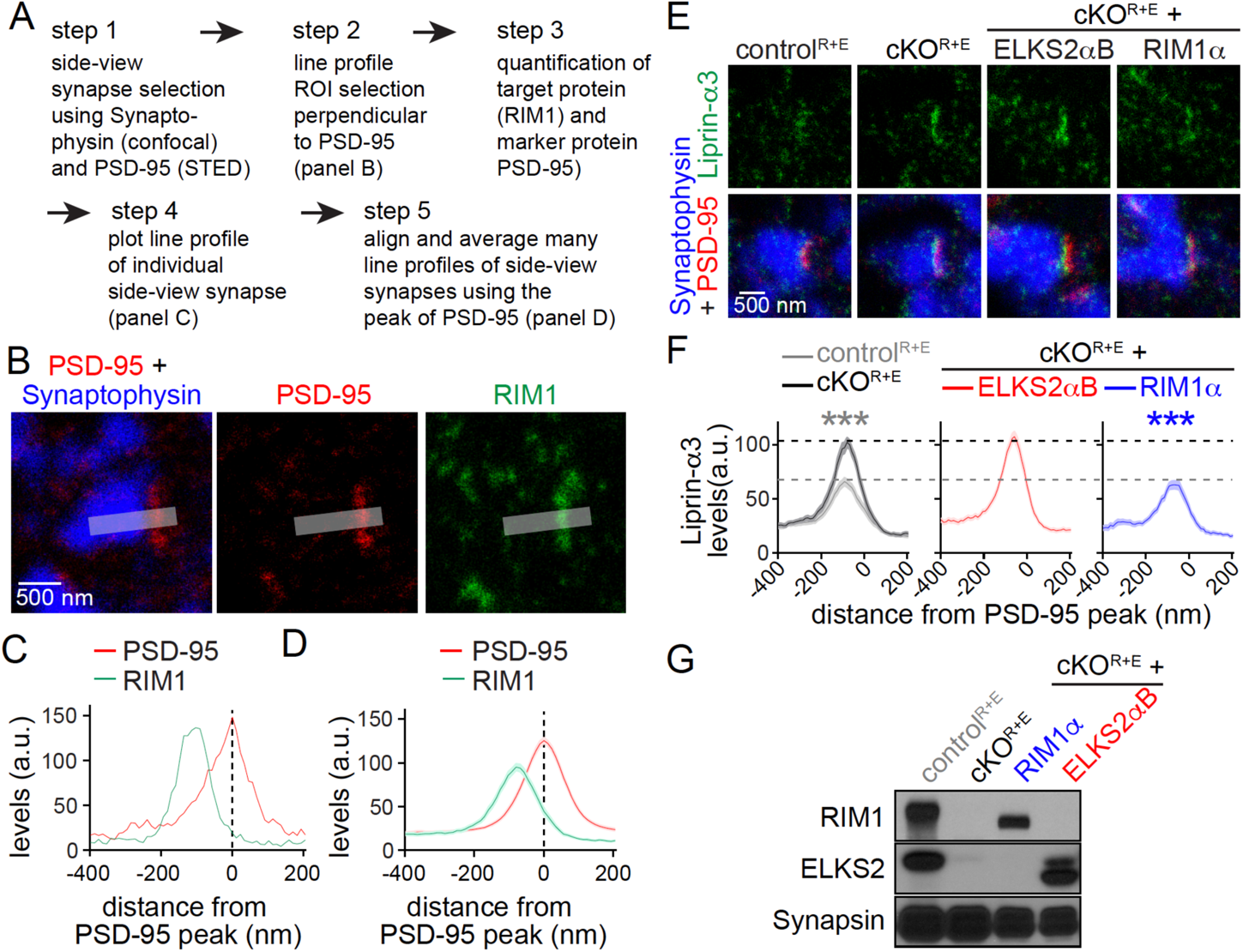
Workflow for STED analyses and assessment of protein expression and localization after rescue of active zone disruption, related to Fig. 1. (**A**) Workflow for STED side-view synapse analyses as described in (Held et al., 2020; Nyitrai et al., 2020). (**B**) Example STED images of a side-view synapse (the control^R+E^ images shown in Fig. 1B are reproduced here) stained for PSD-95 (STED), RIM1 (STED) and Synaptophysin (confocal). (**C**)Quantification of fluorescent signals of PSD-95 and RIM1 labeling of the synapse shown in B. (**D**) Summary data of a full experiment for PSD-95 and RIM1, these are the same data as the control^R+E^ data shown in Fig. 1C. (**E, F**) Sample STED images (E) and quantification (F) of side-view synapses stained for Liprin-α3 (STED), PSD-95 (STED), and Synaptophysin (confocal) of control^R+E^ and cKO^R+E^ neurons, and of cKO^R+E^ neurons after re-expression of ELKS2αB or RIM1α, dotted lines mark control^R+E^ (grey) and cKO^R+E^ (black) levels for comparison, n = 60 synapses/3 independent cultures per condition. (**G**) Western blot for RIM1, ELKS2 and Synapsin in homogenates of control^R+E^ and cKO^R+E^ neurons, and of cKO^R+E^ neurons after re-expression of ELKS2αB or RIM1α. Rescue proteins were expressed at overall levels similar to or below wild type proteins. Data are mean ± SEM; \*\*\**P* < 0.001 compared to cKO^R+E^ as determined by two-way ANOVA followed by Dunnett’s multiple comparisons post-hoc tests.

**Figure S2.**
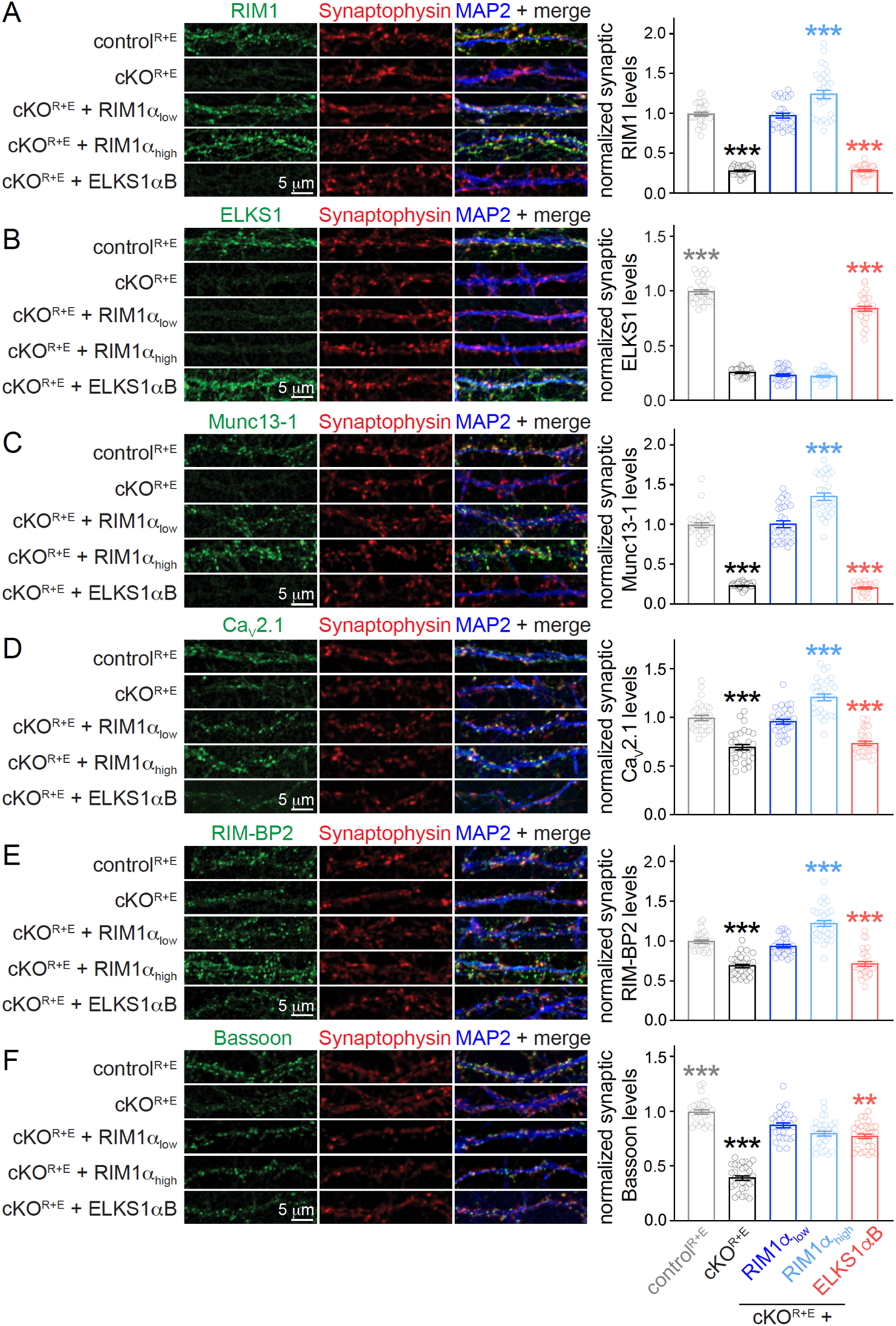
Synaptic RIM1α levels determine levels of interacting active zone proteins, related to Fig. 1. (**A**) Sample confocal images (left) and quantification (right) of RIM1 levels in control^R+E^ and cKO^R+E^ synapses, and in cKO^R+E^ synapses expressing HA-tagged full-length RIM1α or ELKS1αB. Neurons were infected at DIV3 with lentiviruses expressing RIM1α under a synapsin promoter for lower expression (RIM1α_low_), or under a ubiquitin promoter for higher expression (RIM1α_high_). Synaptophysin staining was used to define ROIs, and levels of RIM1 were quantified within those ROIs and normalized in each culture to levels in control^R+E^ neurons. n = 30 images/3 cultures per condition. (**B**-**F**) Same as A, but for ELKS1 (B), Munc13-1 (C), Ca_V_2.1 (D), RIM-BP2 (E), and Bassoon (F). Note that the levels of Munc13-1 (C), Ca_V_2.1 (D), and RIM-BP2 (E) correlate well with levels of RIM (A). Furthermore, in contrast to rescue ELKS2*α*B which does not localize to cKO^R+E^ synapses (Fig. 1), rescue ELKS1*α*B is localized at least in part to synapses and recovers some synaptic Bassoon (F). This may be related to their distinct functions and differential localization, with ELKS2αB at the active zone and ELKS1αB broadly distributed throughout the nerve terminal (Nyitrai et al., 2020). n = 30/3 each. Data are mean ± SEM; \*\**P* < 0.01, \*\*\**P* < 0.001 compared to cKO^R+E^ + RIM1α_low_ as determined by Brown-Forsythe ANOVA followed by Dunnett’s T3 test (A, B, C and E), or by one-way ANOVA followed by Dunnett’s tests (D and F).

**Figure S3.**
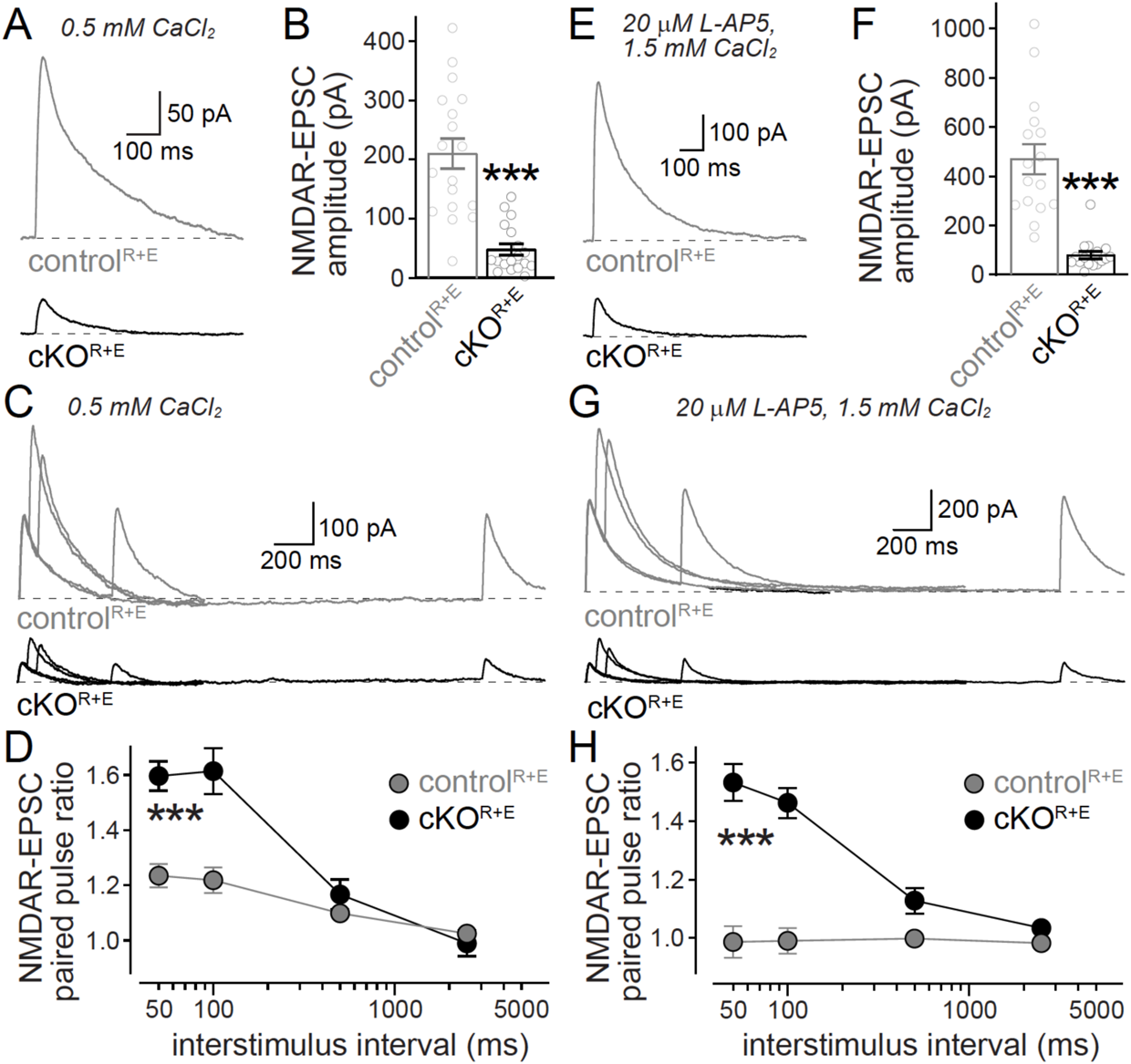
Characterization of NMDAR-mediated synaptic transmission, related to Fig. 2. (**A, B**) Sample traces (A) and quantification (B) of NMDAR-mediated EPSCs evoked by focal electrical stimulation with 0.5 mM Ca^2+^ in the extracellular solution (instead of 1.5 mM used in all other experiments), n = 18 cells/3 independent cultures per condition. (**C, D**) Sample traces (C) and quantification (D) of NMDAR-mediated paired pulse ratios induced with focal electrical stimulation in extracellular solution containing 0.5 mM Ca^2+^ (instead of 1.5 mM used in all other experiments). The relationship of paired pulse ratios between cKO^R+E^ and control^R+E^ was similar in 0.5 (D) or 1.5 mM (Fig. 2H) Ca^2+^, n = 16/3 per condition. (**E, F**) Sample traces (E) and quantification (F) of NMDAR-mediated EPSCs evoked by focal electrical stimulation in 1.5 mM Ca^2+^ and in the presence of 20 µM of the low affinity NMDAR antagonist L-AP5 to prevent NMDAR saturation, n = 16/3 per condition. (**G, H**) Sample traces (G) and quantification (H) of NMDAR-mediated paired pulse ratios induced with focal electrical stimulation in extracellular solution containing 1.5 mM Ca^2+^ and 20 µM L-AP5. Paired pulse ratios between cKO^R+E^ and control^R+E^ were similar with (H) or without (Fig. 2H) L-AP5, n = 16/3 per condition. Overall, A-G indicate that NMDAR saturation does not confound the estimates of p in our standard conditions (1.5 mM Ca^2+^, no L-AP5). Data are mean ± SEM; \*\*\**P* < 0.001 compared to control^R+E^ as determined by Mann-Whitney tests (B and F) or two-way ANOVA followed by Dunnett’s tests (D and H).

**Figure S4.**
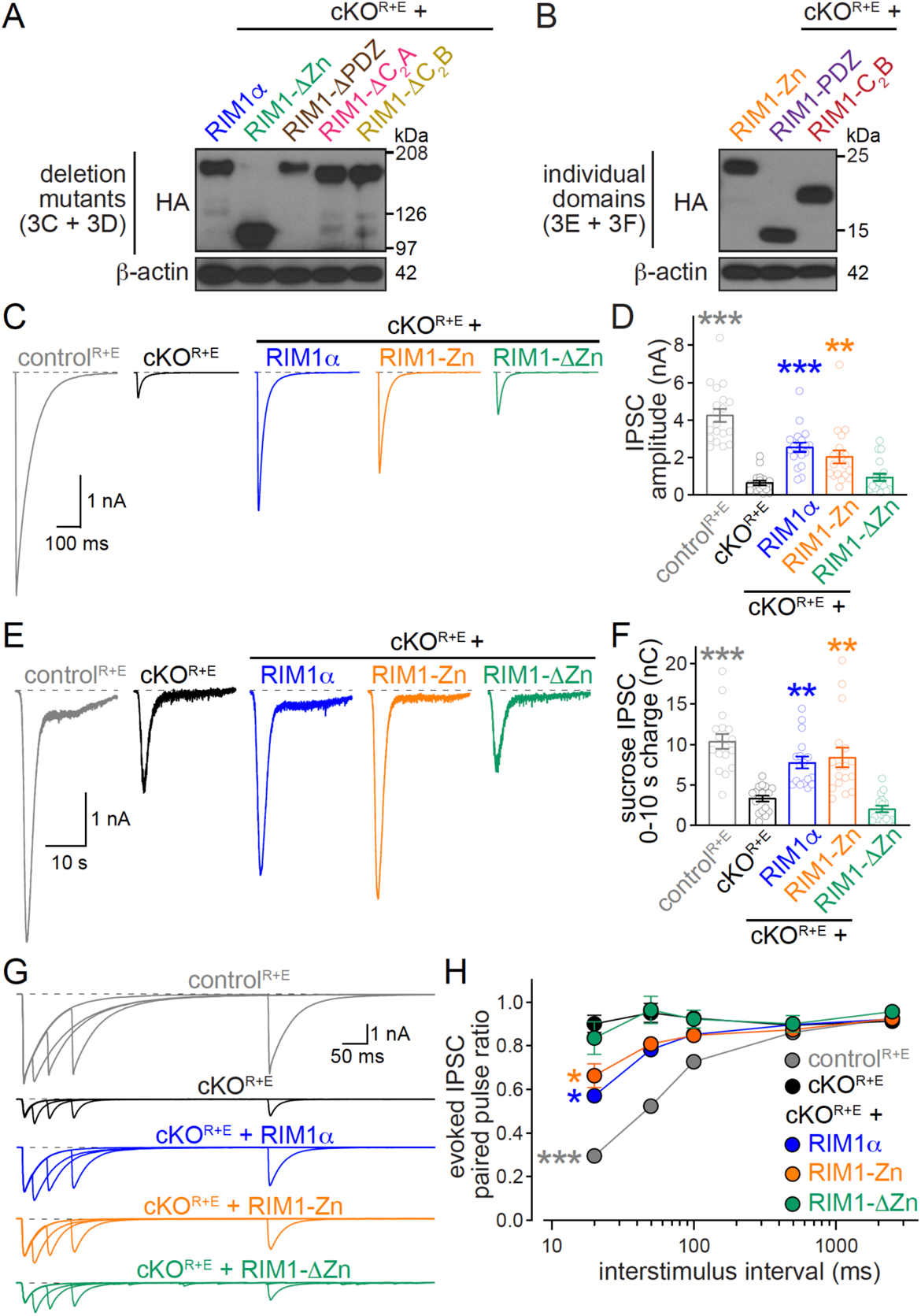
RIM1 zinc finger re-expression restores the RRP at inhibitory cKO^R+E^ synapses, related to figs. 3 and 4. (**A, B**) Expression levels of rescue constructs in homogenates of cultured cKO^R+E^ neurons were assessed with anti-HA antibodies, and anti-β-actin antibodies were used as loading controls. Several RIM1-C_2_A constructs were generated in addition to the constructs that are shown, but their expression could never be detected by Western blotting, and constructs were hence not further used for experimentation. (**C, D**) Sample traces (C) and quantification (D) of IPSCs evoked by focal electrical stimulation, control^R+E^ n = 19 cells/5 independent cultures, cKO^R+E^ n = 19/5, cKO^R+E^ + RIM1α n = 19/5, cKO^R+E^ + RIM1-Zn n = 18/5, and cKO^R+E^ + RIM1-ΔZn n = 19/5. (**E, F**) Sample traces (E) and quantification (F) of IPSC triggered by hypertonic sucrose, the first 10 s of the IPSC were quantified to estimate the RRP, n = 17/3 per condition. (**G, H**) Sample traces (G) and quantification (H) of IPSC paired pulse ratios, control^R+E^ n = 17/5, cKO^R+E^ n = 17/5, cKO^R+E^ + RIM1α n = 18/5, cKO^R+E^ + RIM1-Zn n = 17/5, and cKO^R+E^ + RIM1- ΔZn n = 18/5. Data are mean ± SEM; \**P* < 0.05, \*\**P* < 0.01, \*\*\**P* < 0.001 compared to cKO^R+E^ as determined by Kruskal-Wallis followed by Dunn’s tests (D and F) or two-way ANOVA followed by Dunnett’s tests (H).

**Figure S5.**
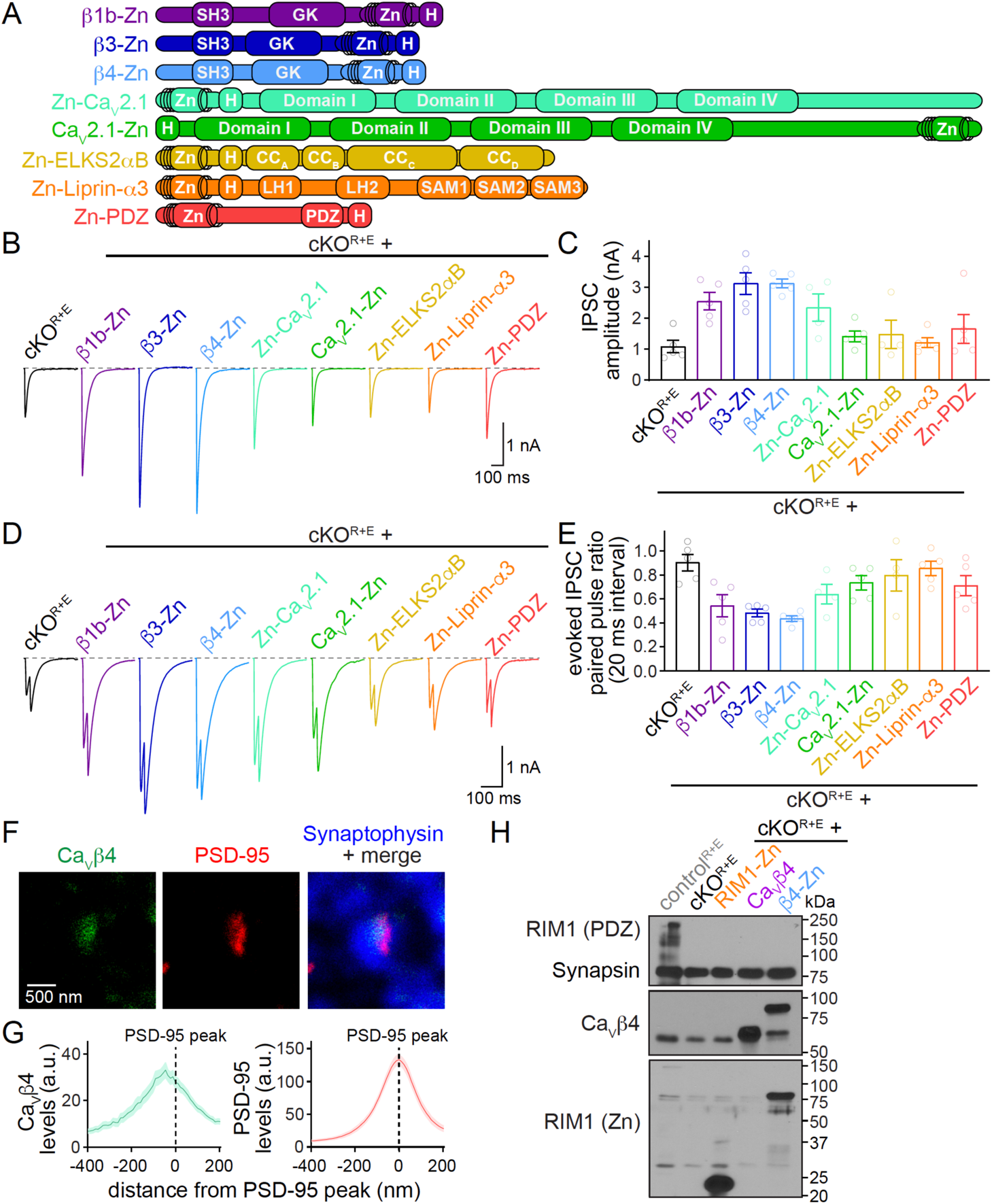
Screening of fusion-proteins for restoring action potential-triggered release at cKO^R+E^ synapses, related to Fig. 5. (**A**) Schematic representation of candidate fusion-proteins of the RIM1 zinc finger domain to Ca_V_β1b, Ca_V_β3, Ca_V_β4, Ca_V_2.1 (N- or C-terminal fusions), ELKS2αB, Liprin-α3 or RIM1-PDZ. (**B, C**) Sample traces (B) and quantification of IPSC amplitudes (C) evoked by a focal stimulation, n = 4-5 cells/1 culture per condition. (**D**, **E**) Sample traces (D) and quantification (E) of IPSC paired pulse ratios at 20 ms interstimulus intervals, n = 4-5 cells/1 culture per condition. (**F, G**) Sample STED images (F) and quantification (G) of side-view synapses stained for Ca_V_β4 (STED), PSD-95 (STED), and Synaptophysin (confocal) in wild type synapses, n = 23 synapses/1 culture. (**H**) Expression levels of RIM1 and rescue constructs in homogenates of cultured control^R+E^ and cKO^R+E^ neurons were assessed with anti-RIM1 PDZ (top), anti-Ca_V_β4 (middle) and anti-RIM1 zinc finger (bottom) antibodies, and anti-Synapsin antibodies were used as loading controls. Data shown as mean ± SEM, no statistics were performed for (C) and (E) due to the limited number of observations. Based on the rescue activity of β4-Zn for evoked IPSC amplitudes and PPR, and the active zone-like localization of endogenous Ca_V_β4, β4-Zn was chosen for full characterization.

**Figure S6.**
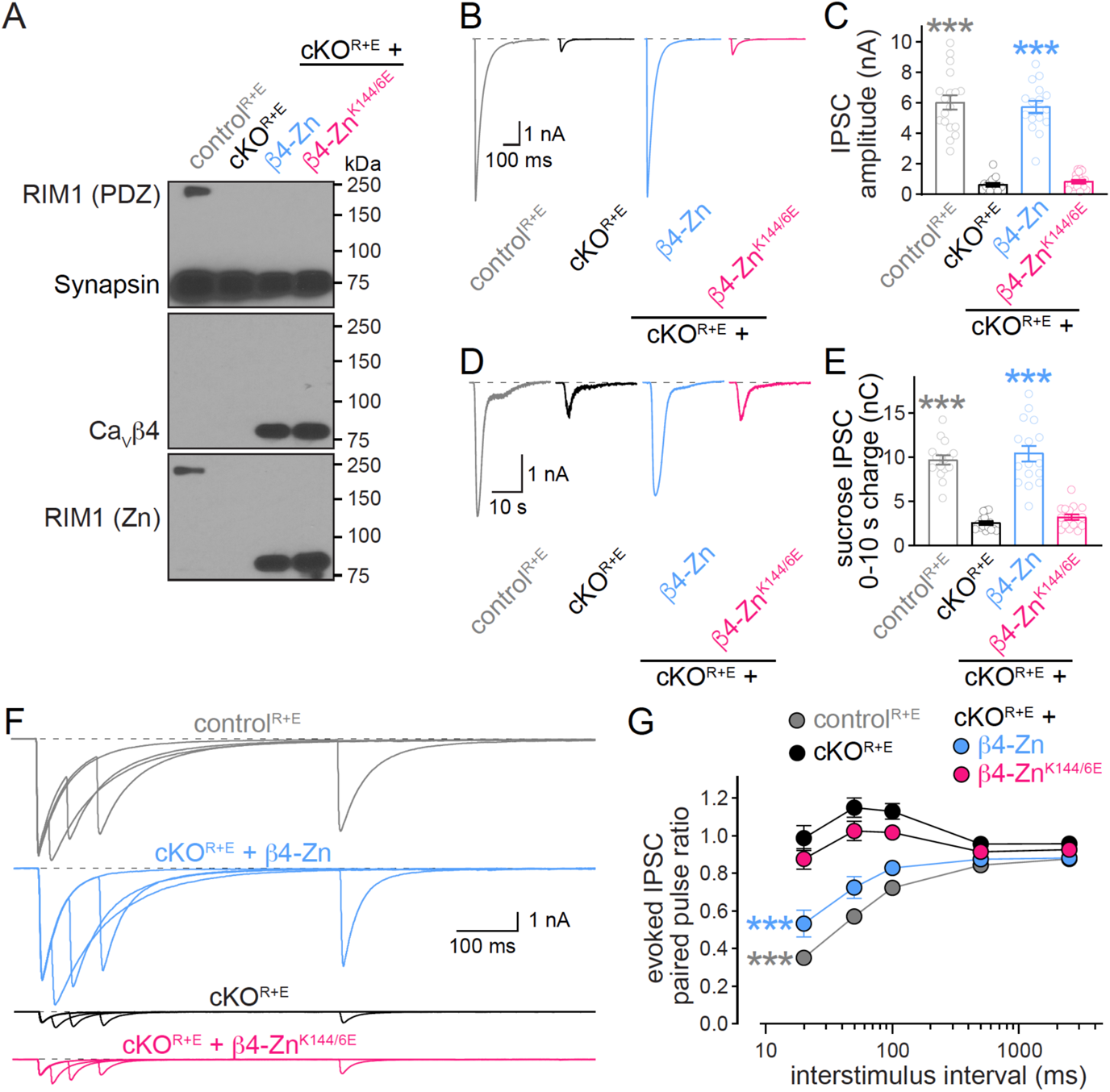
Binding of β4-Zn to Munc13 is essential for restoring release at inhibitory cKO^R+E^ synapses, related to figs. 7 and 8. (**A**) Expression levels of RIM1 and rescue constructs in homogenates of cultured control^R+E^ and cKO^R+E^ neurons were assessed with anti-RIM1 PDZ (top), anti-Ca_V_β4 (middle) and anti-RIM1 zinc finger (bottom) antibodies, and anti-Synapsin antibodies were used as loading controls. (**B, C**) Sample traces (B) and quantification (C) of IPSCs evoked by focal electrical stimulation, control^R+E^ n = 18/3, cKO^R+E^ n = 17/3, cKO^R+E^ + β4-Zn n = 16/3, and cKO^R+E^ + β4-Zn β4-Zn^K144/6E^ n = 18/3. (**D**, **E**) Sample traces (D) and quantification (E) of IPSC triggered by hypertonic sucrose, the first 10 s of the IPSC were quantified to estimate the RRP, n = 16/3 per condition. (**F**, **G**) Sample traces (F) and quantification (G) of IPSC paired pulse ratios, n = 16/3 per condition. Data are mean ± SEM; \*\*\**P* < 0.001 compared to cKO^R+E^ as determined by Kruskal-Wallis followed by Dunn’s tests (C and E) or by two-way ANOVA followed by Dunnett’s tests (G).

